# CRISPR-Cas9-assisted genome editing in *E. coli* elevates the frequency of unintended mutations

**DOI:** 10.1101/2024.03.19.584922

**Authors:** Karl A. Widney, Dong-Dong Yang, Leo M. Rusch, Shelley D. Copley

**Affiliations:** Department of Molecular, Cellular and Developmental Biology, University of Colorado Boulder, Boulder, CO, 80309, USA; Department of Biochemistry, University of Colorado Boulder, Boulder, CO, 80309, USA; Cooperative Institute for Research in Environmental Sciences, University of Colorado Boulder, Boulder, CO, 80205, USA

## Abstract

Cas-assisted lambda Red recombineering techniques have rapidly become a mainstay of bacterial genome editing. Such techniques have been used to construct both individual mutants and massive libraries to assess the effects of genomic changes. We have found that a commonly used Cas9-assisted editing method results in unintended mutations elsewhere in the genome in 26% of edited clones. The unintended mutations are frequently found over 200 kb from the intended edit site and even over 10 kb from potential off-target sites. We attribute the high frequency of unintended mutations to error-prone polymerases expressed in response to dsDNA breaks introduced at the edit site. Most unintended mutations occur in regulatory or coding regions and thus may have phenotypic effects. Our findings highlight the risks associated with genome editing techniques involving dsDNA breaks in *E. coli* and likely other bacteria and emphasize the importance of sequencing the genomes of edited cells to ensure the absence of unintended mutations.

**GRAPHICAL ABSTRACT:** 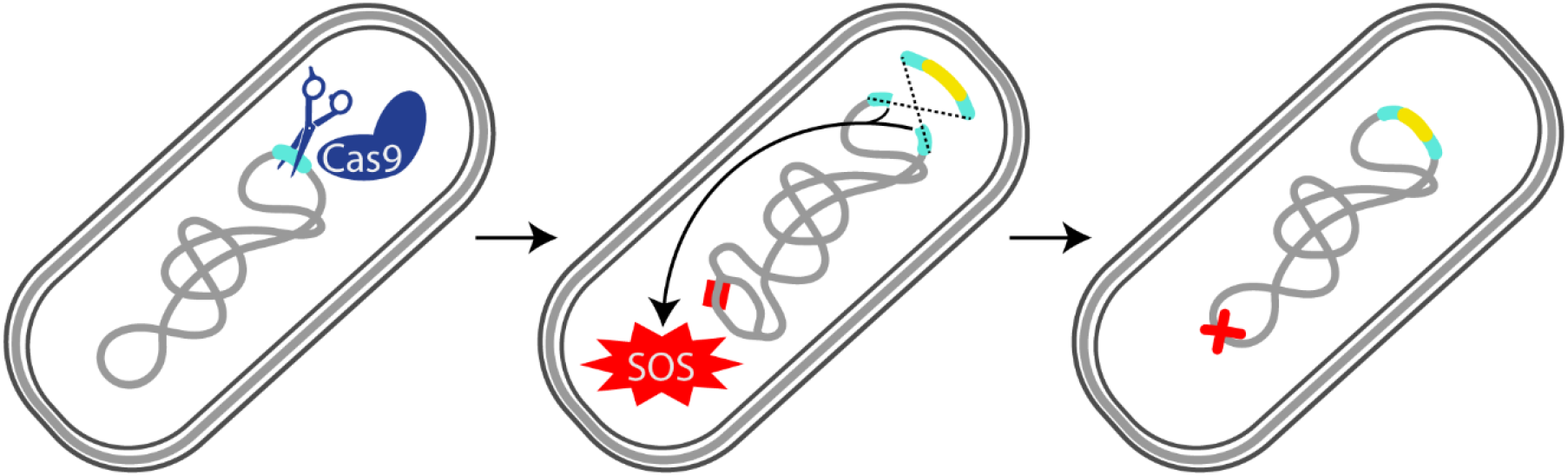

## INTRODUCTION

The type II CRISPR/Cas system from *Streptococcus pyogens* has been adapted for efficient genome editing in many bacteria (1–4). The Cas9 nuclease is directed by a 20-nucleotide spacer sequence in a single guide RNA (sgRNA) to introduce a double-strand DNA break (DSB) in the genome at a complementary “proto-spacer” sequence next to a proto-spacer adjacent motif (PAM) (5, 6). DSBs introduced by Cas9 are generally lethal in bacteria due to the lack of robust repair pathways (7, 8). However, lambda Red-promoted recombination with an editing cassette provided as single-strand (ss) DNA, linear double-strand (ds) DNA or plasmid DNA can heal the DSB with concomitant incorporation of a desired edit (4, 9, 10). Editing cassettes are designed to change or remove the Cas9 target site so as to prevent re-cleavage of the genome after editing. Thus, Cas9 acts as a programmable counter-selection marker that kills unedited cells. Because one mismatch between the spacer and proto-spacer is often insufficient to abolish cutting (11–20), multiple synonymous mutations near the target mutation are often introduced during editing (10, 21). However, synonymous mutations can cause phenotypic effects (22–30). Consequently, we prefer a two-step scarless editing method (24, 31) in which the synonymous “immunizing” mutations introduced during the first step are reverted in the second step, leaving only the desired mutation(s).

Cas9-assisted editing in bacteria has been used to assess the phenotypic effects of individual point mutations (32) and of more extensive edits such as modification of ribosomes (33), introduction of genes encoding new metabolic pathways (34–38) and deletion of virulence genes (1, 39). Additionally, creation of large mutant libraries has enabled screening of the effects of edits on antibiotic resistance (21) and essential gene functions (24, 40). While the introduction of desired edits is typically confirmed by Sanger sequencing, the genomes of edited strains are often not sequenced (1, 3, 32, 35–39). Thus, the possibility that edited cells might harbor unintended mutations elsewhere in the genome is often overlooked.

We routinely use a variety of genome editing procedures [*Streptococcus pyogens* (*Sp*) Cas9-assisted editing (9, 10, 21, 24, 38, 41), lambda Red recombineering (42, 43) and I-SceI-assisted editing (44)] in various strains of *E. coli* as well as other bacteria. We sequence the genomes of edited strains to confirm expected genotypes prior to carrying out phenotypic analyses. Whole-genome sequencing of a large number of edited *E. coli* strains created by two researchers for various projects in the lab revealed unintended mutations in 26% of colonies after a single step of Cas9-assisted editing and after I-SceI-assisted editing. The frequency of colonies with unintended mutations is well above the 10% found in a control experiment as well as the 8% found after lambda Red recombineering. The increase in unintended mutations after editing is not due to off-target activities of Cas9 and I-SceI but is likely a consequence of expression of error-prone polymerases due to introduction of DSBs in the genome.

## MATERIAL AND METHODS

### Biological Resources

Strains and plasmids used in this work are listed in Tables S1 and S2, respectively. Primers are listed in Table S3.

### Cas9-assisted genome editing

All strains subjected to Cas9-assisted editing were derived from *E. coli* MG1655*, a strain of MG1655 with five mutations that had been introduced for a previous project (24), carrying a previously constructed helper plasmid (24) (pDY118A, Figure S1, Addgene ID 182958). pDY118A encodes SsrA-tagged *Sp*Cas9 (10) under control of an anhydrotetracycline (aTc)-inducible promoter, the lambda Red proteins Exo, Beta and Gam under control of a heat-inducible promoter and I-SceI under control of an isopropyl ß-D-1-thiogalactopyranoside (IPTG)-inducible promoter. (The SsrA tag ensures that any Cas9 produced by leaky expression is quickly degraded (10).) Mutations were introduced using editing cassettes carried on donor plasmids (Table S4) or provided as ssDNA (Table S5) (Figure 1A). sgRNAs containing 20-nucleotide (nt) spacer sequences to direct Cas9 cleavage were encoded on donor or guide plasmids constructed from pAM041 (Figure S2, Addgene ID 217969) as described below. (Guide plasmids have backbones identical to those of donor plasmids but lack an editing cassette.) Spacer sequences used to direct Cas9 genome cleavage and editing cassettes are listed in Tables S4 and S5.

**Figure 1.**
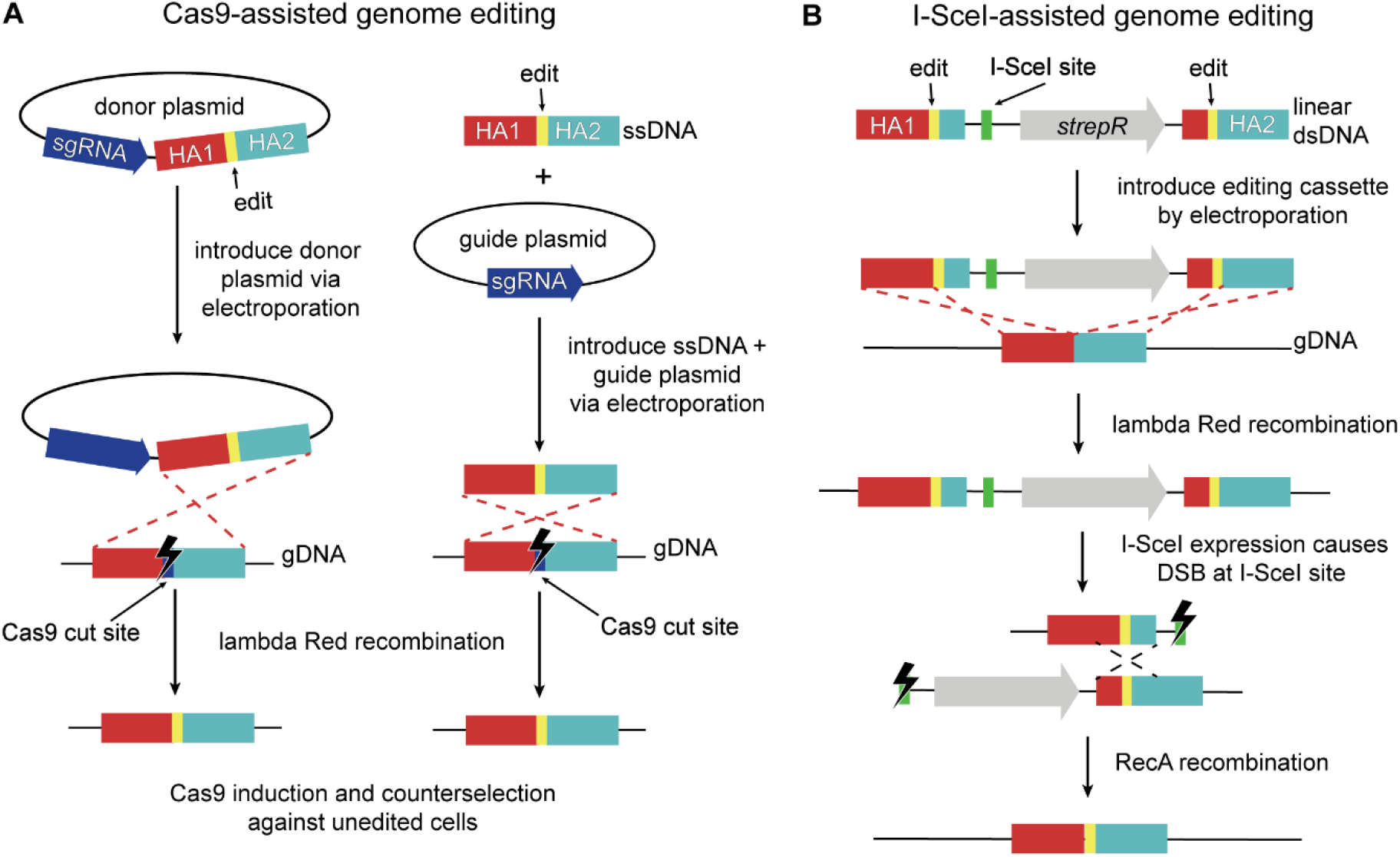
Methods used for Cas9- and I-SceI-assisted genome editing of *E. coli* strains carrying pDY118A, a helper plasmid that encodes Cas9 under control of an aTc-inducible promoter, the lambda Red proteins under control of a heat-inducible promoter, I-SceI under control of an IPTG-inducible promoter and a chloramphenicol resistance gene. Lambda Red genes were induced by heat shock prior to the start of each protocol. HA – homology arm. Figures not drawn to scale. (**A**) Cas9-assisted genome editing uses either donor plasmids that encode both an sgRNA and an editing cassette (left) or guide plasmids encoding an sgRNA along with a 120-nt ssDNA editing cassette (right). Successfully edited cells are selected on LB plates containing ampicillin (100 µg/ml) to select for cells with the donor or guide plasmid, aTc (0.4 µg/ml) to induce expression of Cas9 and chloramphenicol (34 µg/ml) to maintain the helper plasmid. (**B**) I-SceI-assisted genome editing (44) uses editing cassettes introduced as linear dsDNA fragments. Successful recombinants are isolated on LB plates containing streptomycin (100 µg/ml) and chloramphenicol (30 µg/ml). Induction of I-SceI expression with 1 mM IPTG results in a DSB that is healed by RecA-mediated repair, leading to incorporation of the desired edit in most cells (depending upon the point at which homologous recombination occurs).

In preparation for genome editing, cells were streaked from a frozen glycerol stock onto LB agar plates containing 34 μg/ml chloramphenicol (to maintain pDY118A) and grown overnight at 30 ºC. A single colony was inoculated into 5 ml of LB containing 34 μg/ml chloramphenicol and grown overnight at 30 ºC with shaking. Five hundred µl of this culture was inoculated into 50 ml of LB containing 34 μg/ml chloramphenicol. The cells were grown at 30 ºC with shaking to mid-log phase (OD600 ≈ 0.3), transferred to a shaking 42 ºC water bath and incubated for 15 min to induce expression of the lambda Red genes. The cells were then cooled on ice for at least 15 min and prepared for electroporation as described previously (24). A donor plasmid (100 ng) or a guide plasmid (100 ng) and linear ssDNA editing cassette (100 ng) were introduced into washed competent cells (90 µl) by electroporation. The cells were added to 1 ml Super Optimal broth with Catabolite (SOC) repression medium (45), allowed to recover at 30 ºC for 3 h and then plated on fresh LB agar plates containing 34 μg/ml chloramphenicol to maintain pDY118A, 0.4 µg/ml aTc to induce expression of Cas9 and 100 μg/ml ampicillin to select for cells carrying the donor or guide plasmid. The following day, colonies were streaked onto plates containing 34 μg/ml chloramphenicol and 100 μg/ml ampicillin and grown overnight at 30 ºC to eliminate unedited cells from colonies that were mixtures of edited and unedited cells. (Editing is sometimes completed after plating, leading to the presence of unedited cells in some colonies.) Colony PCR using the manufacturer’s protocol for NEB OneTaq Hot Start DNA Polymerase (M0481L) and primers flanking the edited site (Table S3) was used to amplify fragments encompassing intended edit sites. Amplicons were treated with ThermoFisher ExoSap-IT Express (75001) and subjected to Sanger sequencing to verify successful editing. Edited colonies were inoculated into 5 ml of LB. Cultures were grown overnight at 37 ºC with shaking prior to the isolation of genomic DNA (gDNA). Strains generated by Cas9-assisted editing are listed in Dataset S1 – Cas9 edited strains.

### Lambda Red Recombineering

Lambda Red recombineering was used to replace *pdxB* in *E. coli* BW25113, a K12 strain closely related to the MG1655 strain (46, 47) used for Cas9-assisted genome editing, with a kanamycin resistance gene. (The minor differences between the BW25113 and MG1655 strains (https://bioinfo.ccs.usherbrooke.ca/BW25113.html) should not impact editing efficiency or mutagenesis.) *pdxB* was replaced with *kanR* in *E. coli* BW25113 carrying pDY118A using a donor plasmid (pKAW086, Figure S3, Table S2, Addgene ID 217967) containing an editing cassette comprised of *kanR* flanked by regions of homology to the genome (Table S6). pKAW086 cannot be maintained in *E. coli* lacking an exogenous *pir* gene (48), allowing selection of recombinants in which *kanR* has been incorporated into the genome on plates containing kanamycin. The backbone of pKAW086 also contains a gene encoding GFP to enable screening for cells that integrated the entire plasmid into the genome instead of replacing *pdxB* with *kanR*. Alternatively, *pdxB* was replaced with *kanR* using a linear dsDNA editing cassette amplified from pKAW086 using NEB Q5 Hot Start DNA Polymerase and primers listed in Table S3. We found no difference in the frequency of unintended mutations after using plasmid and linear dsDNA editing cassettes and therefore present their results together.

In preparation for genome editing, approximately 10 μl of frozen cells was inoculated into 5 ml of LB containing 30 μg/ml chloramphenicol and grown overnight at 30 ºC with shaking. Fifty µl of this culture was inoculated into 5 ml of LB containing 30 μg/ml chloramphenicol. The cells were grown at 30 ºC with shaking to mid-log phase (OD600 ≈ 0.2), transferred to a shaking 42 ºC water bath and incubated for 15 min to induce expression of the lambda Red genes. The cells were then cooled on ice for at least 15 min and prepared for electroporation as described previously (24). pKAW086 (100 ng) or the linear dsDNA editing cassette (100 ng) was introduced into washed competent cells (50 µl) by electroporation. The cells were added to 1 ml SOC (45), allowed to recover at 30 ºC for 3 h and then plated on LB agar plates containing 30 μg/ml chloramphenicol to maintain pDY118A and 50 μg/ml kanamycin to select for edited cells. The following day, colonies were streaked onto plates containing 30 μg/ml chloramphenicol and 50 μg/ml kanamycin and grown overnight at 30 ºC to eliminate unedited cells from colonies that were mixtures of edited and unedited cells. Individual colonies were inoculated into 10 µl of phosphate-buffered saline (PBS). One µl of the colony resuspension was subjected to colony PCR using the manufacturer’s protocol for NEB OneTaq Hot Start DNA Polymerase (M0481L) and primers flanking the edited site (Table S3). Amplicons were treated with ThermoFisher ExoSap-IT Express (75001) and subjected to Sanger sequencing to verify successful replacement of *pdxB* with *kanR*. Five µl of the colony resuspensions was inoculated into 5 ml of LB. Cultures were grown overnight at 37 ºC with shaking prior to isolation of gDNA. Strains generated by lambda Red recombineering are listed in Dataset S1 – lambda Red only strains.

### I-SceI-assisted genome editing

Edits were introduced into Δ*pdxB*::*kan E. coli* BW25113 (BW25113*, generated as described above) containing pDY118A or strains derived from BW25113* containing pDY118A by previous rounds of editing using a three-part dsDNA editing cassette comprised of: first, a segment containing a homology arm (HA) upstream of the edit, the edit and a short downstream HA; second, a segment containing an 18-bp I-SceI cut site and a streptomycin resistance gene; and third, a final segment containing a short HA upstream of the edit, the edit and a downstream HA (Figure 1B). The editing cassette (Table S7) was amplified from pKAW066 (Figure S4, Table S2, Addgene ID 217968) using the manufacturer’s protocol for NEB Q5 Hot Start DNA Polymerase (M0494L) and primers listed in Table S3.

Lambda Red proteins were expressed and cells were prepared for electroporation as described for lambda Red recombineering. The editing cassette (100 ng) was introduced into washed competent cells (50 µl) by electroporation, after which the cells were added to 1 ml SOC at 30 ºC and allowed to recover for 3 h and then plated on LB agar plates containing 30 μg/ml chloramphenicol to maintain pDY118A and 50 μg/ml streptomycin to select for cells that had successfully recombined the editing cassette into the genome. After overnight incubation at 30 ºC, colonies were streaked onto LB agar plates containing 30 μg/ml chloramphenicol and 50 μg/ml streptomycin and grown overnight at 30 ºC to ensure elimination of unedited cells. Individual colonies were then streaked onto fresh LB agar plates containing 30 μg/ml chloramphenicol and 1 mM IPTG to induce I-SceI expression and promote RecA-mediated recombination to obtain the scarless edit (Figure 1B). After overnight incubation at 30 ºC, colonies were screened for incorporation of the desired edit as described for lambda Red editing. Strains generated by I-SceI-assisted editing are listed in Dataset S1 – I-SceI edited strains.

### Whole-genome sequencing and analysis

Genomic DNA (gDNA) was purified using Invitrogen PureLink gDNA kits (K182002) or NEB Monarch gDNA kits (T3010). DNA concentrations were measured using an Invitrogen Qubit 2.0 fluorometer with the Broad Range dsDNA assay (Q33265). gDNA samples were sent to SeqCenter or SeqCoast for Illumina short-read sequencing. gDNA samples were sometimes pooled at defined percentages (typically 18, 32 and 50%) so that mutations could be ascribed to individual clones based on the abundance of each mutation. This approach decreases the per-clone cost of whole-genome sequencing while still providing adequate coverage (generally >20x per clone) to enable identification of mutations. Raw reads were cleaned and sorted with Fastp 0.23.2 (49) using settings to deduplicate reads, trim the last base on each read and remove reads with an overall quality score of less than 15. The reads were then analyzed using Breseq 0.35.4 (50) and aligned to the MG1655 genome (GenBank U00096.3 (47)) or the BW25113 genome (GenBank CP009273.1 (46)) as well as to the helper plasmid pDY118A and the donor/guide backbone of pAM041 using settings to analyze reads as a population, only show mutations in more than 10% of the reads aligning to a region, and skip bowtie2-stage2 (a stage that aligns fewer than 0.1 % reads and adds unnecessary time to the analysis). (Mapping reads to the plasmid reference files prevented erroneous assignment of plasmid reads to the genome, which could be misinterpreted as mutations in the genome.) The mutations identified by Breseq were exported to a CSV file and sorted in Excel; known mutations in the parental strains and intended mutations created by genome editing that were not in the reference genomes were identified and removed to yield a final list of unintended mutations.

### Prediction of off-target sites and comparison to the locations of unintended mutations

Downloadable versions of Cas-OFFinder 2.4.1 (14) (https://github.com/snugel/cas-offinder) and Cas-OFFinder bulge (https://github.com/hyugel/cas-offinder-bulge) were used to create lists of potential off-target sites for the spacer sequences used in this study. These lists were filtered to exclude sites with more than a total of six mismatches *plus* gaps and more than two gaps. Distances between unintended mutations and the closest potential off-target sites were calculated.

### Plasmid construction

The vector backbones for the donor, guide and control plasmids were amplified from pAM041 using the manufacturer’s protocol for NEB Q5 Hot Start DNA Polymerase (M0494L) and primers listed in Table S3. After PCR, the reaction mixtures were subjected to gel electrophoresis (1% agarose) and bands of the correct size were extracted with a ThermoFisher GeneJET gel extraction kit (K0691) to remove primer dimers. Extracted DNA was treated with NEB DpnI (R0176L) to remove template DNA before being purified using a Monarch PCR & DNA cleanup kit (T1030L).

The empty donor plasmid pKAW109 (containing no editing cassette or sgRNA, Figure S5) was prepared from the amplified vector backbone by treatment with NEB T4 polynucleotide kinase (M0201S) to phosphorylate the 5’ ends of the DNA and NEB T4 ligase (M0202S) to circularize the DNA.

Guide and donor plasmids were prepared by Gibson assembly of the vector backbone and an additional fragment containing an sgRNA and, in the case of donor plasmids, an editing cassette. For guide plasmids, complementary 60-nt oligos (Table S8) containing a spacer sequence and 20-nt regions of homology to the guide plasmid backbone on either end of the spacer sequence were purchased from IDT and annealed by boiling the two oligos in PBS and allowing them to cool slowly to 28 ºC. For donor plasmids, gBlocks (Table S4) containing editing cassettes and spacer sequences were obtained from IDT. Insert and vector backbone fragments were mixed with NEB Gibson Assembly Master Mix (E2611S) and incubated for 1-4 hours at 50 ºC. The reaction mixtures were dialyzed at room temperature for 1-2 h against distilled water using MF-Millipore 0.025 µm MCE membranes (VSWP02500) and used to transform electrocompetent *E. coli* DH5α cells as previously described (24). The cells were allowed to recover in SOC (45) at 37 ºC for one hour and then plated on LB containing 100 μg/ml ampicillin. Correct assembly was confirmed by colony PCR as described for Cas9-assisted editing with primers spanning the assembly junctions (Table S3). A colony containing correctly assembled plasmid was inoculated into 5 ml LB containing 100 μg/ml ampicillin and grown overnight with shaking at 37 ºC. Plasmids were purified from overnight cultures using a Monarch® Plasmid Miniprep Kit (T1010L).

The plasmids used in lambda Red recombineering and I-SceI-assisted editing were also constructed by Gibson assembly (51) and have been deposited to AddGene.

### Reagents

Chemicals (antibiotics, IPTG, etc.) and media (LB and SOC) were purchased from Sigma-Aldrich and Thermo-Fischer. Enzymes used for plasmid construction were purchased from NEB. Primers were obtained from IDT and Eurofins genomics. gBlocks were obtained from IDT. A homemade Gibson master mix was made according to the recipe on dx.doi.org/10.17504/protocols.io.n9xdh7n.

### Statistical Analyses

A multiple proportions Chi-square test was performed in Excel using a spreadsheet available at (https://www.moresteam.com/university/downloads/calculators.xls). Adjusted p-values for pairwise comparisons were obtained using the Marascuilo procedure (52).

### Data Availability

All data underlying this article are available in the article and its online supplementary material.

## RESULTS

### Unintended mutations occur frequently after Cas9-assisted genome editing

We used Cas9-assisted genome editing techniques similar to established literature protocols (9, 10, 21, 24, 38, 41) (Figure 1A) to introduce point mutations into the genomes of strains derived from *E. coli* MG1655* (24), a strain of MG1655 with five mutations constructed for a previous project. We utilized the helper plasmid pDY118A (24), which encodes Cas9 under control of an aTc-inducible promoter, lambda Red proteins under control of a heat-inducible promoter and the homing endonuclease I-SceI under control of an IPTG-inducible promoter. (Although I-SceI was encoded on the helper plasmid, expression was not induced during Cas9-assisted editing.) We edited loci in eight different genes in strains derived from MG1655* for various projects in the laboratory using editing cassettes introduced either on a donor plasmid or as linear ssDNA (Tables S4 and S5). In total, we sequenced the genomes of 117 colonies after Cas9-assisted editing. The intended edit was successfully introduced in all cases. However, unintended mutations introduced by genome editing (i.e. mutations that were not present in the precursor strain) were found in 26% of the colonies; 29 had one unintended mutation and two had two unintended mutations (Figure 2, Dataset S1 – Cas9 edited strains). All but two of the unintended mutations were unique; the two identical mutations may have arisen in the population prior to editing. Notably, the unintended mutations were found as far as 2.2 Mb from the intended edit.

**Figure 2.**
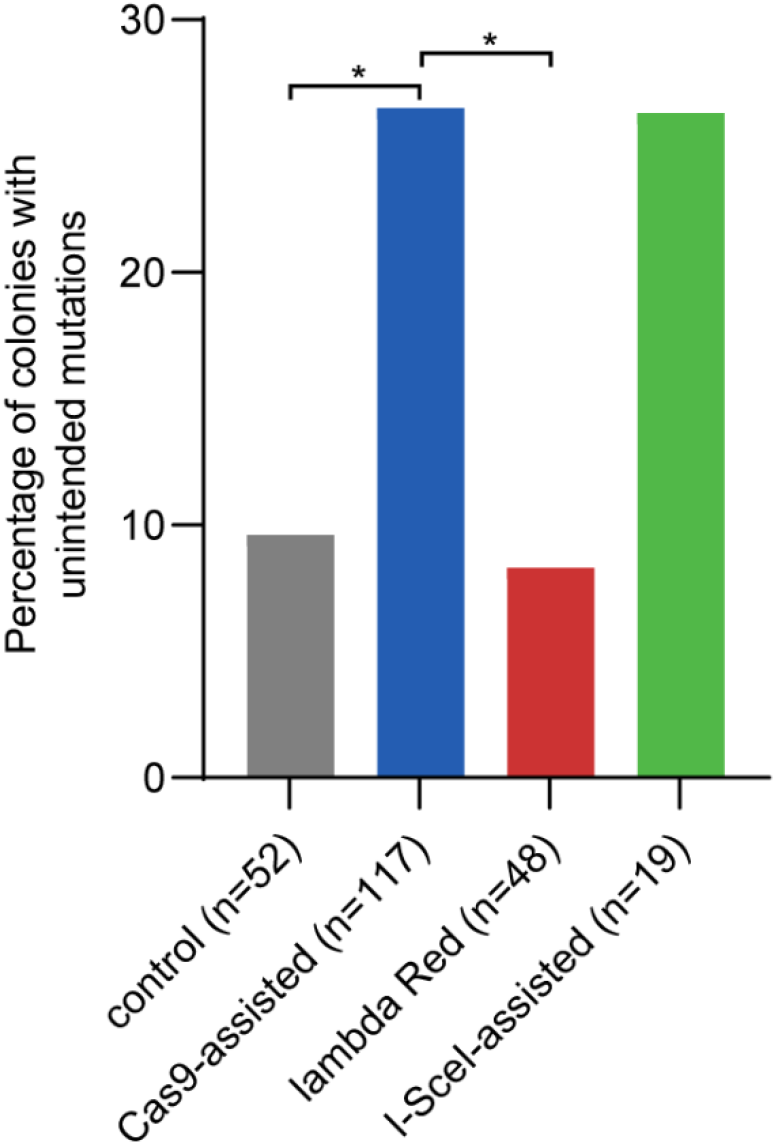
The percentage of colonies with unintended mutations produced by three different editing protocols and one control protocol. *, adjusted p < 0.05 using the Marascuilo procedure (52) for comparing multiple proportions. All other comparisons are not statistically significant.

Some unintended mutations in edited clones would be expected due to the intrinsic rate of mutations introduced during genome replication or as a result of error-prone DNA damage repair. In our Cas9-assisted protocol, editing was initiated with a single colony. By the time editing was completed, the cells had undergone 70-80 cell divisions. To determine the frequency of unintended mutations arising during an equivalent number of cell divisions without genome editing, we carried out a control experiment using *E. coli* BW25113 (a K12 strain of *E. coli* that is closely related to MG1655 (46, 47)) carrying the helper plasmid pDY118A and an empty donor plasmid (Figure S5). Lambda Red proteins and Cas9 were expressed but unable to perform editing due to the lack of an editing cassette and sgRNA in the empty donor plasmid. We sequenced the genomes of 52 colonies and discovered unique unintended mutations in five (10%); four had one unintended mutation and one had two unintended mutations (Figure 2, Dataset S1 – control strains). Based upon the mutation rate reported for mutation accumulation experiments in *E. coli* MG1655 (2.0 x 10^−10^ per nt per generation, or 9.3 x 10^−4^ per genome per generation) (53), we expect mutations in 7-8% of colonies after 70-80 cell divisions, a range close to the value observed for the control experiment. These results suggest that the frequency of unintended mutations after Cas-assisted genome editing is significantly higher than expected from the low background mutation rate.

### The high frequency of unintended mutations after Cas9-assisted genome editing is not due to off-target Cas9 activity

Cas9 is known to have off-target activity that can introduce DSBs in sequences similar to the intended cut site specified by the spacer sequence (11–20). Off-target activity is of particular concern in large eukaryotic genomes (12), but should be less problematic for smaller bacterial genomes because the probability of finding two highly similar 20-nt target sequences in the genome is low. Nevertheless, we considered the possibility that off-target activity might contribute to the high frequency of unintended mutations in our edited clones.

If unintended mutations are due to DSBs at off-target sites, we would expect them to occur in similar places when a common spacer is used (12). We had used a single spacer sequence to create mutations in four different genes in *E. coli* MG1655*. This spacer targets a 23-bp sequence from the gene encoding mCherry that is cleaved efficiently by Cas9. We often use this sequence in a two-step editing protocol that produces a scarless genome edit (24) (i.e. an edit with the desired mutation(s) but no synonymous mutations near the PAM site to prevent re-cleavage of the genome after editing.) In the first step, we introduce a DSB at the editing site using Cas9 and provide an editing cassette that introduces the mCherry sequence. After confirming that the strains produced in the first step have not acquired any unintended mutations, we introduce a DSB in the mCherry sequence using Cas9 and provide an editing cassette that restores the original gene sequence but with the desired mutation(s). Even with the common mCherry spacer sequence, 11 of the 14 unintended mutations found in 63 edited colonies were scattered throughout the genome (Figure 3A). Two of the 14 unintended mutations were identical and may have arisen in the population prior to editing. These two mutations were within 90 bp of another unintended mutation. These three unintended mutations appear as one line on the Circos plot. This region may be a hot-spot for mutations, but it is 35 kb from the nearest potential off-target site predicted by Cas-OFFinder (14). Further, this site is a poor match for the spacer sequence in the sgRNA; it has five mismatches and one gap compared to the target site. Thus, the unintended mutations are unlikely to be due to error-prone repair of DSBs introduced by off-target Cas9 cleavage.

**Figure 3.**
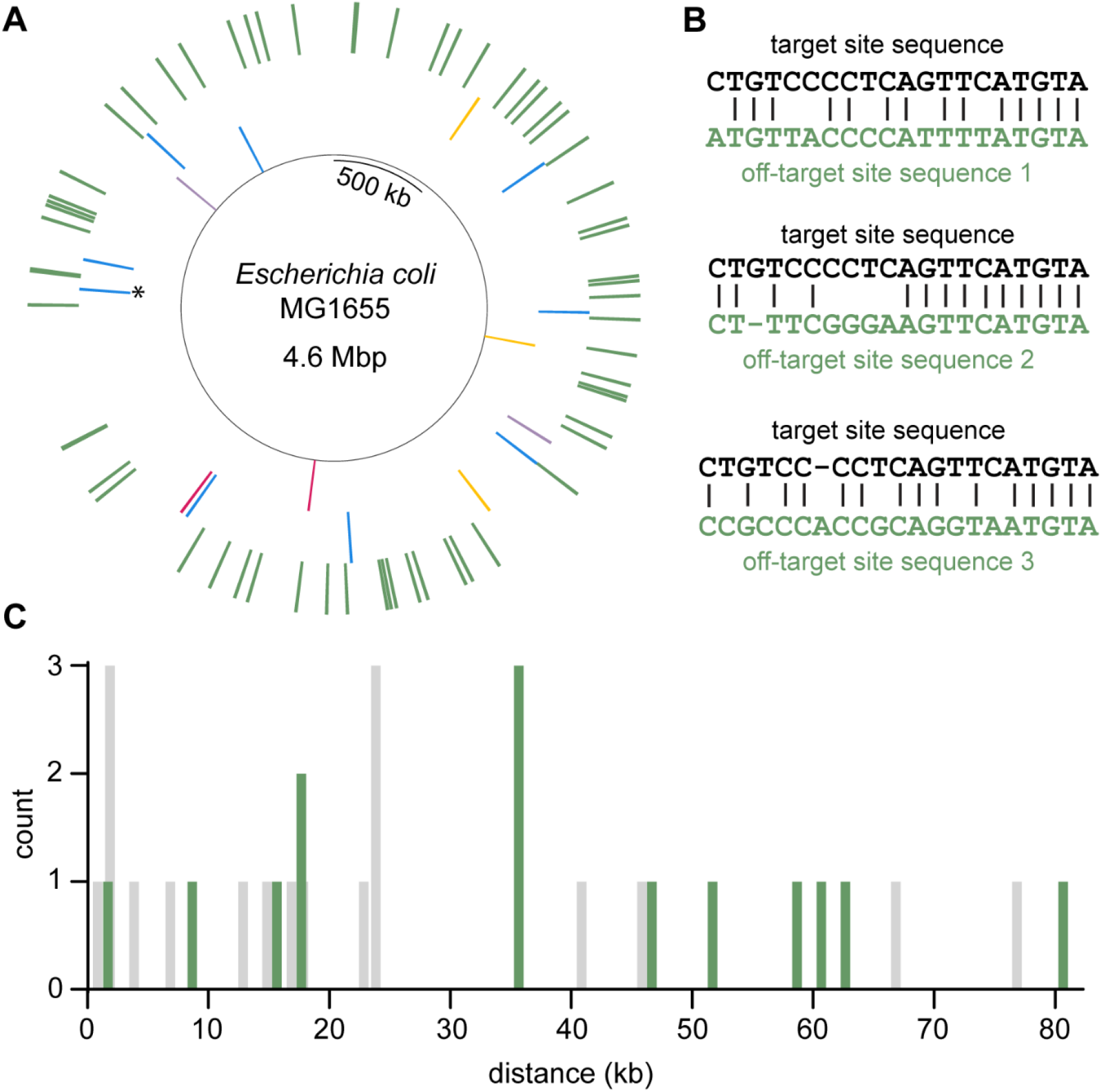
Unintended mutations are not close to predicted off-target sites. (**A**) Circos plot showing the locations of unintended mutations in clones edited with the same sgRNA. This sgRNA targets a 23-bp mCherry sequence introduced during a previous round of genome editing. Inner ring, intended edit sites; second ring, unintended mutations color-coded according to the intended edit site (two edit sites within one kb of each other are both shown in blue); outer ring, off-target sites predicted by Cas-OFFinder with no more than six mismatches plus gaps and a maximum of two gaps. * marks a site where two identical mutations and a third mutation occurred within 90-bp. The plot was generated using the python package PyCircos available at https://github.com/ponnhide/pyCircos. (**B**) Alignments between the target site and three off-target sites predicted by Cas-OFFinder. Sequence one has no gaps and has the first mismatch furthest from the PAM site of off-target sites with no gaps. Sequences two and three have a gap in either the genomic DNA or the sgRNA, and the first mismatch is furthest from the PAM site of off-target sites with a gap in the genomic DNA or sgRNA, respectively. (**C**) Distances between unintended mutations and the closest off-target site for (green) the most-used spacer that targets the mCherry sequence and (grey) the other five spacers. Bin size is one kb and bars are centered around the middle of the bin.

To further probe the possibility that the unintended mutations were due to off-target Cas9 activity, we calculated the distance between each unintended mutation and the closest potential off-target site. We used Cas-OFFinder (14) to generate a list of all sites in the genome with a maximum of six mismatches and a maximum of two gaps compared to the target sequence. We filtered this list to allow a maximum of six mismatches *plus* gaps, parameters that are generally accepted as the limit of Cas9 sequence permissiveness (11, 13, 18, 19). We identified 138 off-target sites (Figure 3A, Dataset S2 – CTGTCCCCTCAGTTCATGTA), all of which were poor matches to the spacer sequence, containing a total of five or more mismatches plus gaps. Three of the best predicted off-target sites are shown in Figure 3B. The distances between unintended mutations and the nearest off-target site are plotted in Figure 3C. Unintended mutations in strains edited with this spacer were an average of 38 kb away from the nearest off-target site.

To ensure that these results were not specific to the spacer that targets the mCherry sequence, we used Cas-OFFinder (14) to predict possible off-target sites for the five other spacer sequences we had used. The other spacers had 78-321 potential off-target sites with six or fewer mismatches plus gaps (Dataset S2). Unintended mutations in strains edited with these five spacers were an average of 22 kb away from the nearest off-target site (Figure 3C).

Only five out of the 33 unintended mutations were found within 2 kb of a potential off-target site. These off-target sites were a poor match for the spacer sequence; four of the five had six mismatches plus gaps, which should prevent Cas9 cleavage. These data demonstrate that unintended mutations cannot be ascribed to error-prone repair in the immediate vicinity of a DSB introduced at either the intended edit site or an off-target site.

### Lambda-Red recombineering does not elevate the frequency of unintended mutations

We considered the possibility that lambda Red recombineering might contribute to the elevated frequency of unintended mutations in our edited strains. We used a standard lambda Red recombineering protocol (42) to replace *pdxB* in *E. coli* BW25113 carrying the helper plasmid pDY118A with an editing cassette encoding kanamycin resistance provided as either dsDNA (Table S6) or on a plasmid (Figure S3). (Expression of genes encoding Cas9 and I-SceI on the helper plasmid was not induced for this protocol.) By the time recombineering was completed, the cells founding colonies picked for whole-genome sequencing had undergone the same number of cell divisions and single-cell-bottlenecks as cells subjected to Cas9-assisted editing. We sequenced 48 colonies and found unique unintended mutations in four (8%); each strain contained only one mutation (Figure 2, Dataset S1 – lambda Red only strains). Thus, our implementation of lambda Red recombineering does not appear to be mutagenic.

### I-SceI-assisted lambda Red recombineering may also elevate the frequency of unintended mutations

We previously developed a scarless genome editing protocol for *E. coli* using the meganuclease I-SceI (44) (Figure 1B) that has largely been superseded by Cas9-assisted genome editing. In the first step of the protocol, an editing cassette containing an antibiotic resistance maker, an 18-bp I-SceI recognition site and regions of homology flanking the desired edit (Figures 1B and S4) is introduced into the genome using lambda Red recombineering. In the second step, induction of I-SceI results in a DSB in the editing cassette. RecA-facilitated recombination between the homology arms in the editing cassette repairs the break, introducing the desired edit in most surviving cells. Therefore, this protocol and the Cas9-assisted protocol share the use of lambda Red and DSB-producing nucleases. However, the I-SceI protocol requires approximately 30 more cell divisions, an additional single-cell bottleneck and relies on both lambda Red- and RecA-mediated recombination.

We used the I-SceI-assisted editing protocol to introduce a single-base pair insertion into *rpoS* in *E. coli* BW25113* and several strains derived from it for other projects in the lab (Table S1). All strains carried the helper plasmid pDY118A. Although Cas9 was encoded on the helper plasmid, it was not used for I-SceI-assisted editing, and any Cas9 produced by leaky expression would have been ineffective due to the lack of an sgRNA. We sequenced the genomes of 19 edited strains and found unique unintended mutations in five (26%); four had one unintended mutation and one had two unintended mutations (Figure 2, Dataset S1 – I-SceI edited strains). All but two of the unintended mutations were unique; the two identical mutations may have arisen in the population prior to editing and were 12.5 kb from the intended edit site. The other four were over 208 kb from the intended edit site. These data suggest that the unintended mutations are not due to error-prone repair in a local region around the DSB introduced during editing.

I-SceI has a known off-target site in the *E. coli* IS5 element (Figure 4A) (54). The IS5 element occurs ten times in the genome of *E. coli* BW25113 (46); two are within 40 kb of each other and appear as a single line in the Circos plot (Figure 4B). Off-target cutting of the genome is likely infrequent since *E. coli* overexpressing I-SceI (albeit from a single genomic copy) has no noticeable growth defects (54). Additionally, unintended mutations that occurred during I-SceI-assisted editing were an average of 312 kb away from the closest off-target site (Dataset S2 – I-SceI sites). These data demonstrate that the unintended mutations in strains after I-SceI editing were not due to off-target activity.

**Figure 4.**
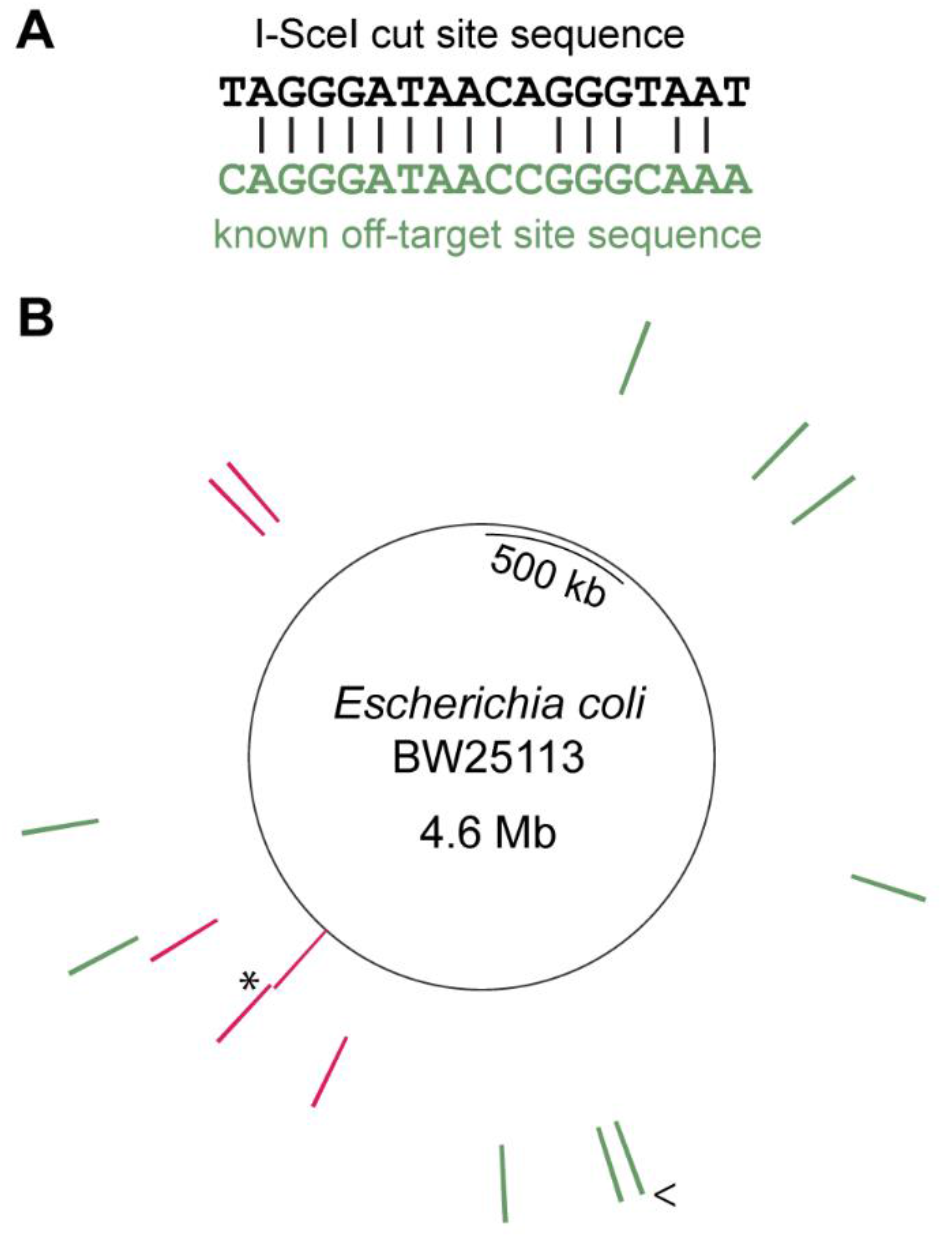
Unintended mutations are not close to known or potential I-SceI off-target sites. (**A**) Alignment of the I-SceI cut site with a known off-target site. (**B**) Circos plot of the BW25113 genome. Inner ring, intended edit site; second ring, unintended mutations; outer ring, known off-target sites in the IS5 element. * marks the site of two identical mutations. < marks a site where two IS5 elements occur within 40 kb of each other. The plot was generated using the python package PyCircos available at https://github.com/ponnhide/pyCircos.

The percentage of colonies containing unintended mutations after I-SceI-assisted editing is the same as that observed after Cas9-assisted editing. However, due to the relatively small sample size, the difference between percentages of unintended mutations in the I-Sce-I-edited strains and the control strain is not statistically significant.

## DISCUSSION

Routine whole-genome sequencing showed that 26% of colonies contained unintended mutations after Cas9-assisted genome editing in *E. coli*, a disturbingly high percentage. A similar elevation of strains containing unintended mutations was observed after I-SceI-assisted genome editing. Only 10% of colonies contained unintended mutations after a control experiment. The increased frequency of unintended mutations is not due to expression of the lambda Red proteins; only 8% of colonies harbored unintended mutations after lambda Red recombineering. These results suggest that Cas9- and I-SceI-assisted genome editing nearly triples the frequency of unintended mutations.

Cas9 and I-SceI both have off-target activities (11–20, 54). Cas9 can generally tolerate up to two mismatches plus gaps if they occur more than seven bp upstream of the PAM site (11–20). Additional mismatches or mismatches closer to the PAM site often lower Cas9 activity, and five or more mismatches usually abolish Cas9 activity (11–20). Our analysis shows that all potential off-target sites for our most-used spacer sequence have five or more mismatches plus gaps, suggesting that off-target activity should be of little concern for this spacer sequence. Further, the unintended mutations found after genome editing were an average of 34 kb away from the closest of these poor off-target sites and, even when the same spacer sequence was used, were spread across the genome. The unintended mutations found after I-SceI-assisted editing were over 50 kb away from known I-SceI off-target sites. These findings establish that the unintended mutations are not likely to be due to error-prone repair after off-target cleavage of the genome by either Cas9 or I-SceI.

Both Cas9- and I-SceI-assisted genome editing introduce DSBs. The I-SceI-assisted protocol necessarily proceeds through a DSB to obtain the final mutation (44). However, Cas9-assisted editing can theoretically occur via two routes. In the first route, lambda Red recombination of the editing cassette into the genome by lambda Red abolishes the Cas9 cut site, preventing DSBs in successfully edited cells. In the second route, Cas9 produced from leaky expression introduces a DSB and then lambda Red recombines the editing cassette into the genome. The observation that the editing rate of Cas9-assisted lambda Red recombineering is significantly higher than that for lambda Red recombineering alone (over 8-fold with some sgRNAs (41)) indicates that the latter route is more common. Thus, most cells going through Cas9-assisted genome editing encounter DSBs.

Previous research has demonstrated that DSBs introduced by Cas9 and I-SceI induce the SOS response (55–57). The SOS response upregulates, among other proteins, three error-prone polymerases (Pol II, IV and V) (58) and increases mutation frequency in undamaged DNA (59) as well as at sites of DNA damage. Pol V is particularly implicated in SOS-induced mutagenesis (60). This enzyme causes base pair substitutions rather than insertions or deletions (61, 62); this mutation spectrum resembles that observed in our edited strains, in which 90% of unintended mutations were single point mutations.

Pol V is a heterotrimer consisting of two UmuD subunits and one UmuC subunit. Activation of the enzyme requires RecA*-facilitated autoproteolytic cleavage of the UmuD subunits to generate UmuD′ as well as association with RecA and ATP to generate a UmuD′2UmuC-RecA-ATP “mutasome” complex (63, 64). (RecA* is the activated form of RecA that forms filaments on ssDNA exposed at a site of DNA damage.) This elaborate mechanism ensures that the enzyme will only be active in the presence of DNA damage so as to avoid unnecessary error-prone repair; due to its lack of 3′-exonuclease proof-reading capability, Pol V introduces errors at a rate of 10^−3^-10^−5^ per base (65), much higher than the 10^−7^ per base error rate (66) of the highly accurate Pol III.

The observation that constitutively expressed Pol V can cause mutations in the absence of DNA damage (67) is likely due to its ability to substitute for Pol III at stalled replication forks, allowing translesion synthesis (68). Additionally, Pol V in association with the β clamp can fill in single-strand gaps left when the Pol III replisome skips over a DNA lesion (60, 69, 70). Pol V mutagenicity via these mechanisms is limited by its poor processivity (71, 72), although it can rebind downstream of a primary repair patch and introduce additional mutations for hundreds of base pairs (60, 72). The unintended mutations we observe are typically found at a considerable distance (up to 2.2 Mb) from the DSBs introduced by Cas9 or I-SceI, so clearly are not caused by a local effect of Pol V at the actual site of the DSB. We suspect that these distant mutations result from the 13-fold increase in the concentration of Pol V due to induction of the SOS response (73), which likely increases the probability that Pol V will substitute for Pol III when the replisome stalls due to an unrepaired lesion or a collision with a transcribing RNA polymerase anywhere in the genome.

The unintended mutations introduced during Cas9- and I-SceI-assisted genome editing are a cause for concern. Most of the 39 unintended mutations we identified were synonymous, nonsynonymous and nonsense mutations point mutations in coding sequences. Among the mutations in intergenic regions, 67 % were within 100 bp of the start codon of a gene and therefore might affect gene expression. Thus, unintended mutations introduced by genome editing could certainly have phenotypic effects that might either augment or diminish the effect of intended mutations.

The occurrence of unintended mutations may be further exacerbated in other genome editing protocols. We found that lambda Red recombineering was not mutagenic using a 15 minute induction of lambda Red expression from genes controlled by the tightly regulated heat-inducible PL promoter and *c*I857^ts^ repressor (74). However, overnight overexpression of lambda Red proteins is mutagenic in *E. coli* (75). Thus, protocols using longer expression times (35, 39) and/or less tightly regulated promoters (9, 35–37, 39), such as the arabinose-inducible promoter (76), may suffer from an elevated frequency of unintended mutations. Protocols that use I-SceI or Cas9 to cure donor/guide plasmids (5, 34, 77–80) may also have elevated mutation frequencies. Such protocols include I-SceI recognition sites or self-targeting sgRNAs on the donor/guide plasmids. DSBs in the donor/guide plasmids efficiently cure the plasmids. However, the additional DSBs generated in the cells may also increase the frequency of unintended mutations.

The problem presented by the unexpectedly high frequency of unintended mutations after genome editing in bacteria might be alleviated in several ways. The frequency of unintended mutations might be decreased by using a version of Cas9-assisted editing that uses a plasmid-encoded dominant negative *recA* mutant to transiently inhibit induction of the SOS response (56). Another promising option is the use of prime editing (81), which avoids introduction of DSBs. However, this approach is currently unsuitable for high-throughput methods and the potential for introducing unintended mutations has not yet been evaluated. Additionally, even the intrinsically low mutation rate of *E. coli* results in unintended mutations in 7-10% of clones after the 70-100 cell divisions required for current genome editing procedures. Our results highlight the importance of verifying that all edited strains are free of unintended mutations by whole-genome sequencing. Although this procedure increases the time and cost required for genome editing, it is essential for accurate assessment of the impact of genomic modifications.

## DATA AVAILABILITY

The data underlying this article are available in the main text and supplementary material. Illumina fastq files, the alignments of reads from edited strains and the gnomes of parental strains are available on NCBI under Bioproject PRJNA1088182 (https://www.ncbi.nlm.nih.gov/sra/PRJNA1088182). Links including Biosample accession numbers are provided for each strain in the supplementary material.

## AUTHOR CONTRIBUTIONS

Karl A. Widney: Conceptualization, Formal analysis, Investigation, Validation, Visualization, Writing—original draft.

Dong-Dong Yong: Conceptualization, Formal analysis, Investigation, Validation, Writing—review & editing.

Leo M. Rusch: Conceptualization, Investigation.

Shelley D. Copley: Conceptualization, Formal analysis, Methodology, Writing—review & editing.

## FUNDING

This work was supported by National Institutes of Health grant numbers R01GM135364 and R01GM124365 to SDC and a Cooperative Institute for Research in Environmental Sciences Graduate Research Fellowship to KAW. Funding for open access charge: National Institutes of Health/ R01GM135364.

## CONFLICT OF INTEREST

The authors declare no conflicts of interest.

## Notes

### Competing Interest Statement

The authors have declared no competing interest.

