## Supplemental figures and tables for "CRISPR-Cas9-assisted genome editing in *E. coli* elevates the frequency of unintended mutations"

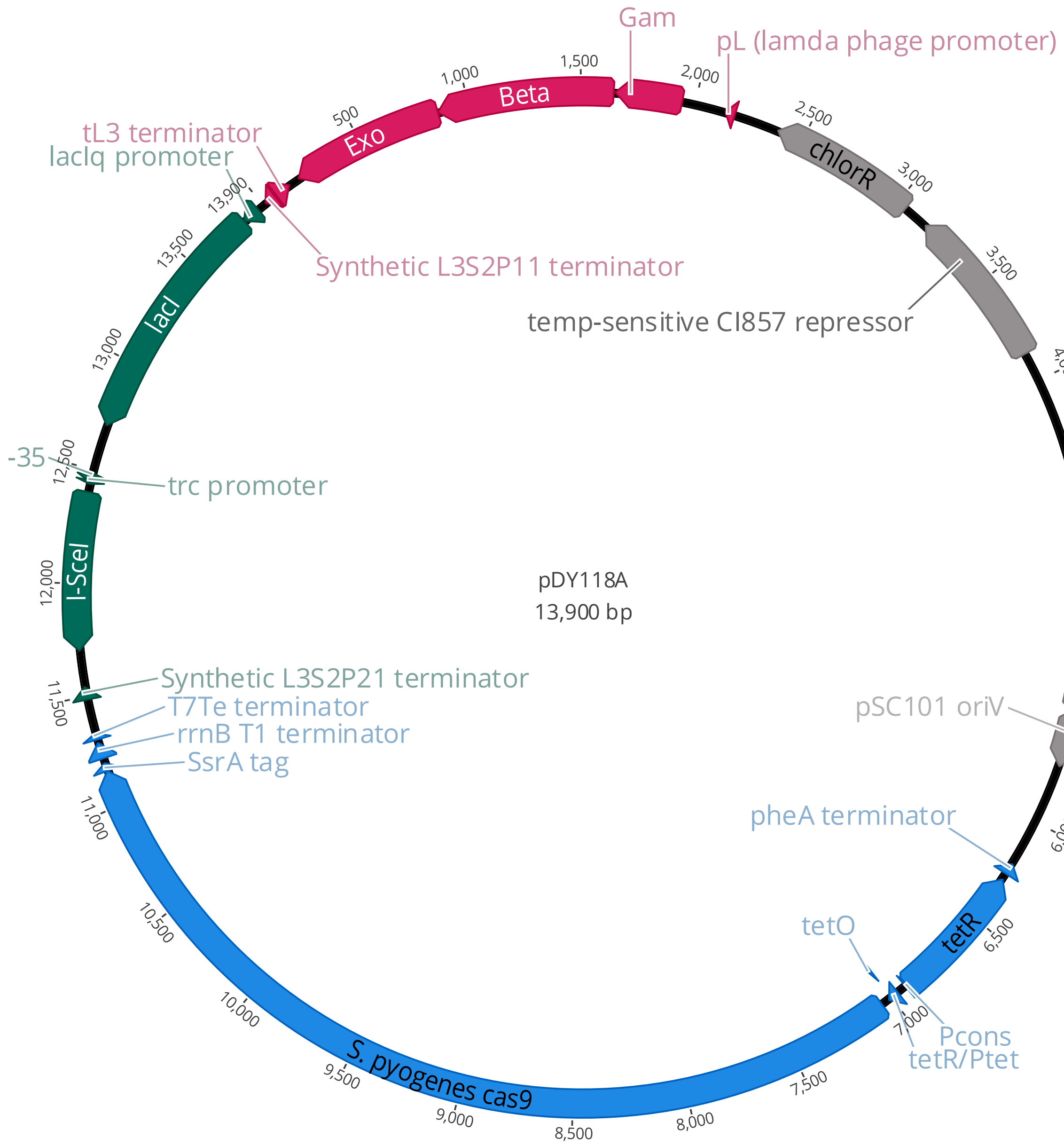


**Figure S1**. Helper plasmid pDY118A. Blue, SsrA-tagged *S. pyogens* Cas9 under regulation of an anhydrotetracycline-inducible promoter; green, I-SceI under regulation of an IPTG-inducible promoter; red, lambda Red proteins Exo, Beta and Gam under regulation of a heat-inducible promoter. pSC101 temperature-sensitive origin of replication allows plasmid to be maintained at 30 °C and cured at 37 °C.

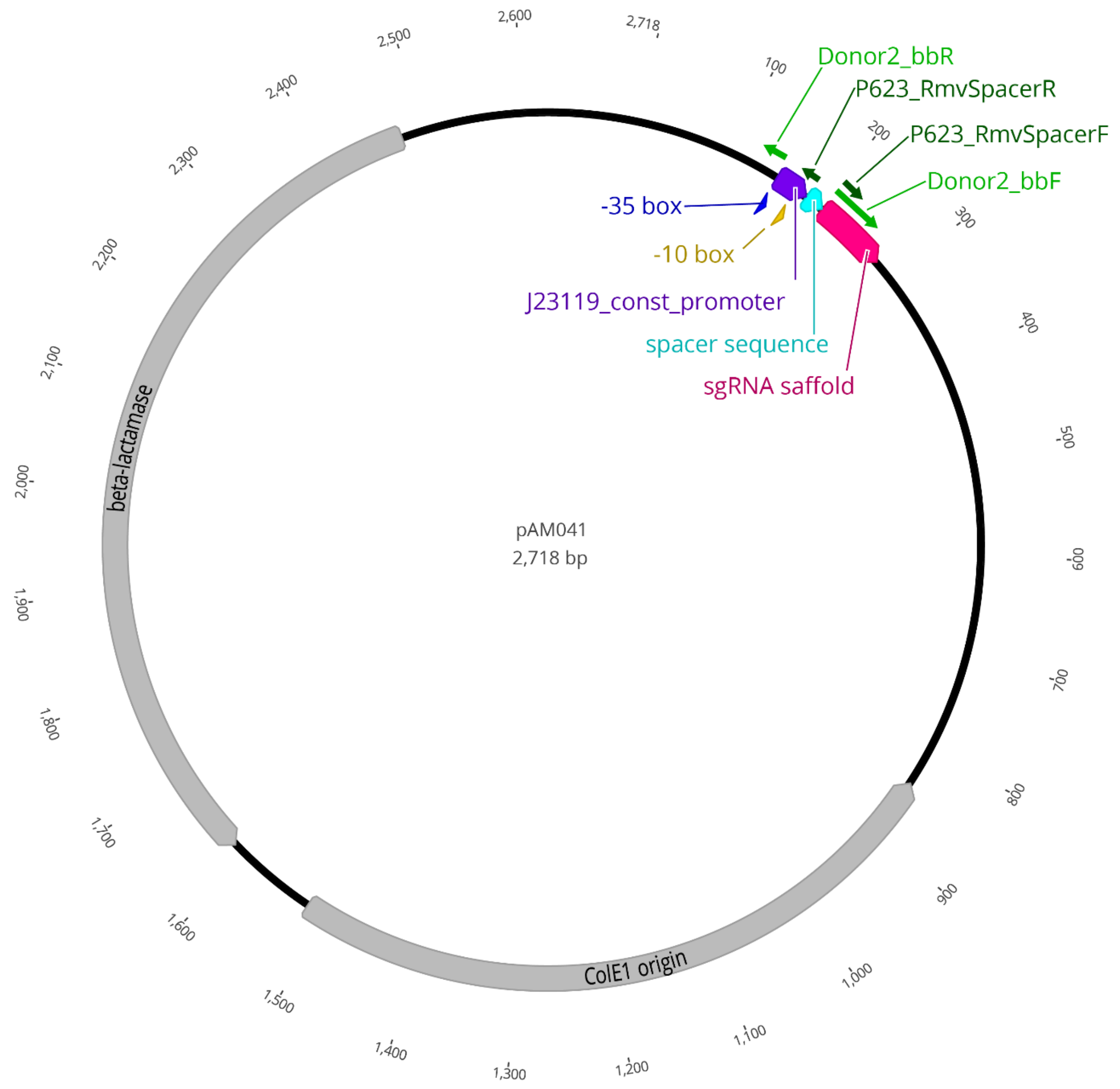


**Figure S2.** pAM041, the plasmid from which backbones for donor and guide plasmids were amplified. Constructed from pSS9_gRNA (1). Purple, promoter for sgRNA; cyan, 20-nucleotide spacer sequence targeting sequences; red, sgRNA scaffold; dark and light green, primer pairs used to amplify the backbone for guide and donor plasmids, respectively. Insertion site for editing cassettes used in donor plasmids is immediately upstream of the promoter.

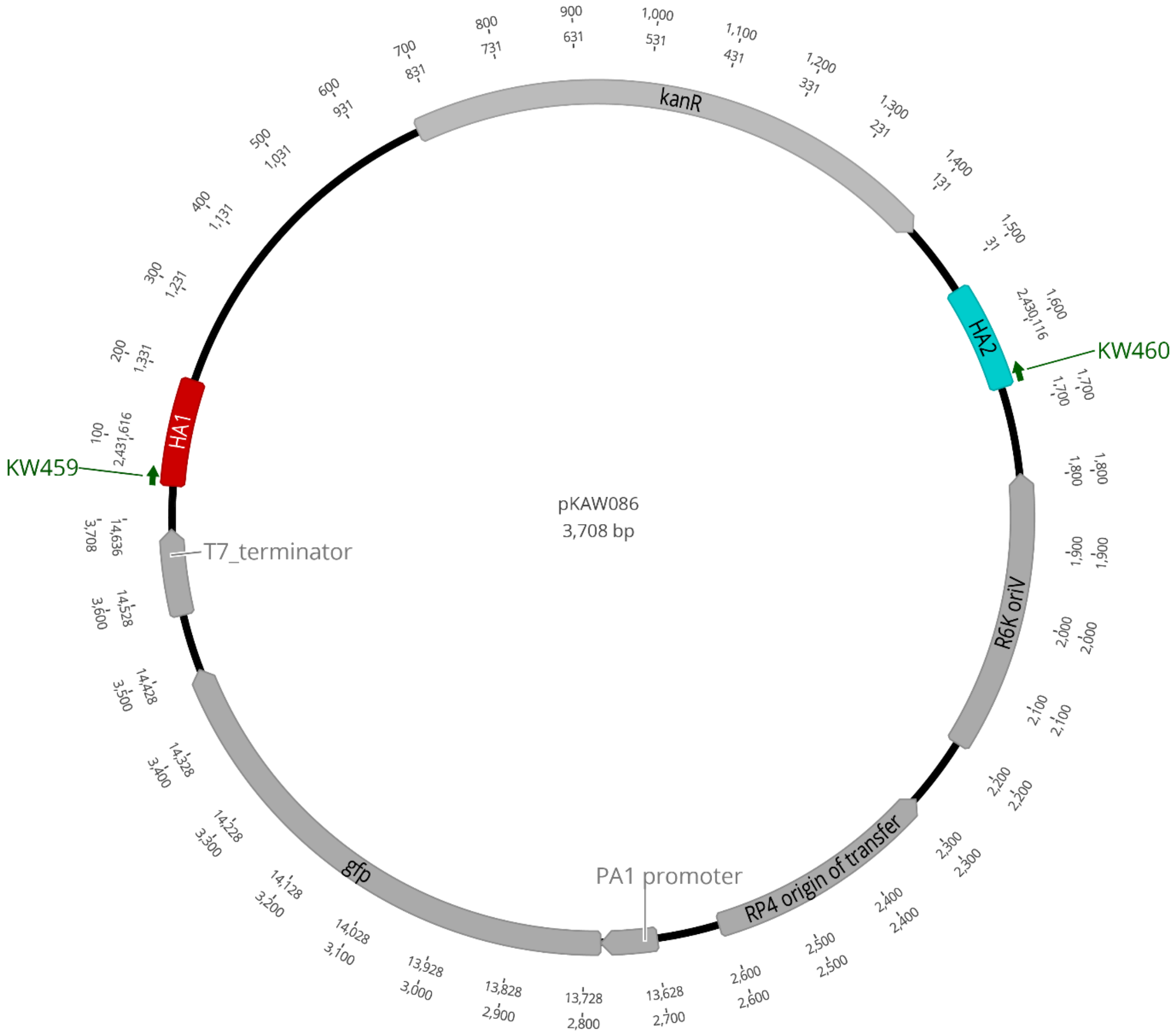


**Figure S3.** pKAW086, which contains the editing cassette used in lambda Red recombineering experiments. Red and cyan, regions with homology upstream and downstream of *E. coli pdxB,* respectively; dark green, primers used to amplify the editing cassette.

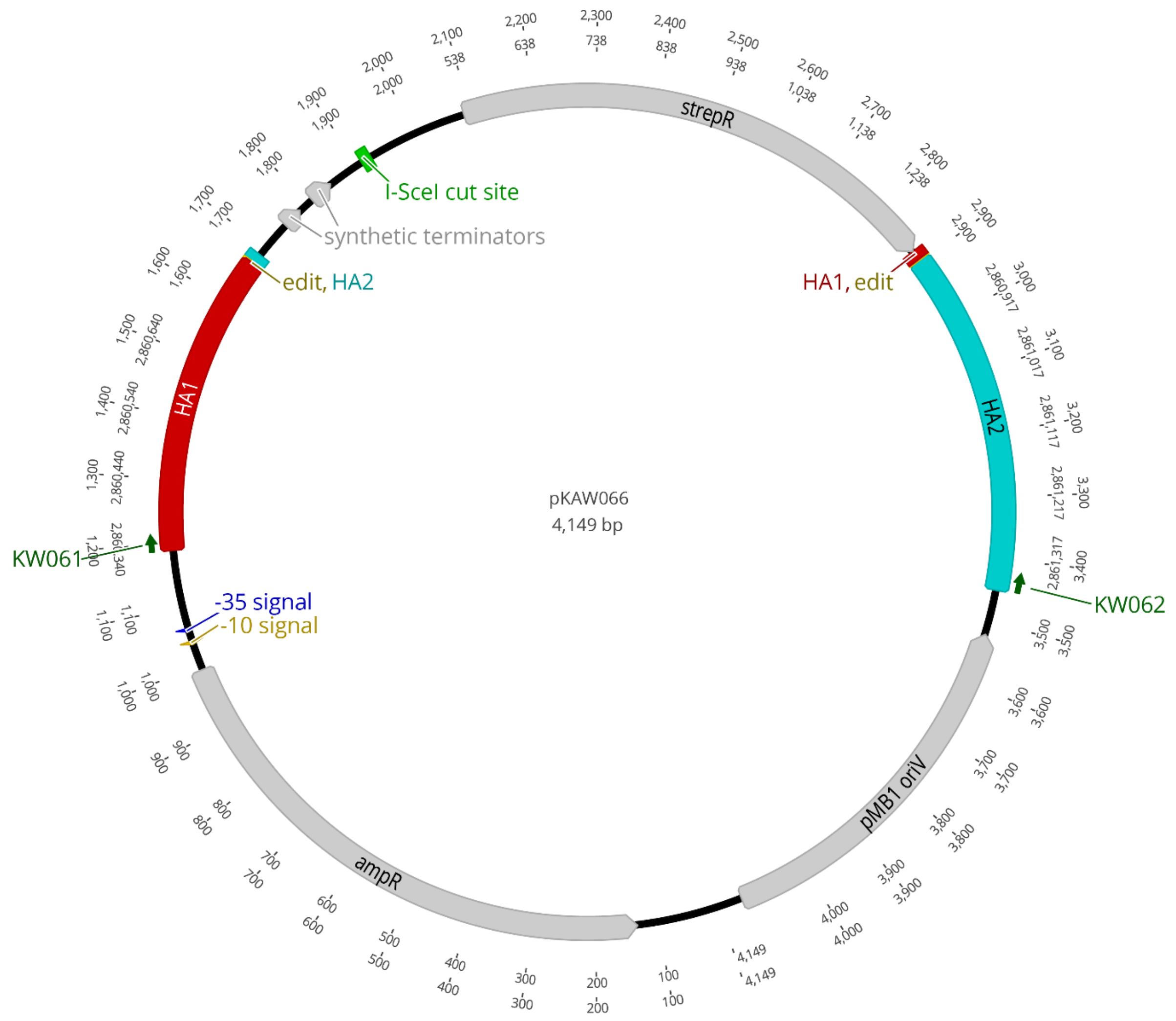


**Figure S4**. pKAW066, which contains the editing cassette used in I-SceI and is comprised of three segments: first, a segment containing a homology arm (HA) upstream of the edit (large red HA1), the edit (yellow) and a short downstream HA (small cyan HA2); second, a segment containing two synthetic terminators (the synthetic terminators ensure efficient cleavage at the I-SceI site), an I-SceI cut-site (green) and a streptomycin resistance gene (grey); and third, a segment containing a short HA upstream of the edit (small red HA1), the edit (yellow) and a downstream HA (large blue HA2). Dark green, primers used to amplify the editing cassette.

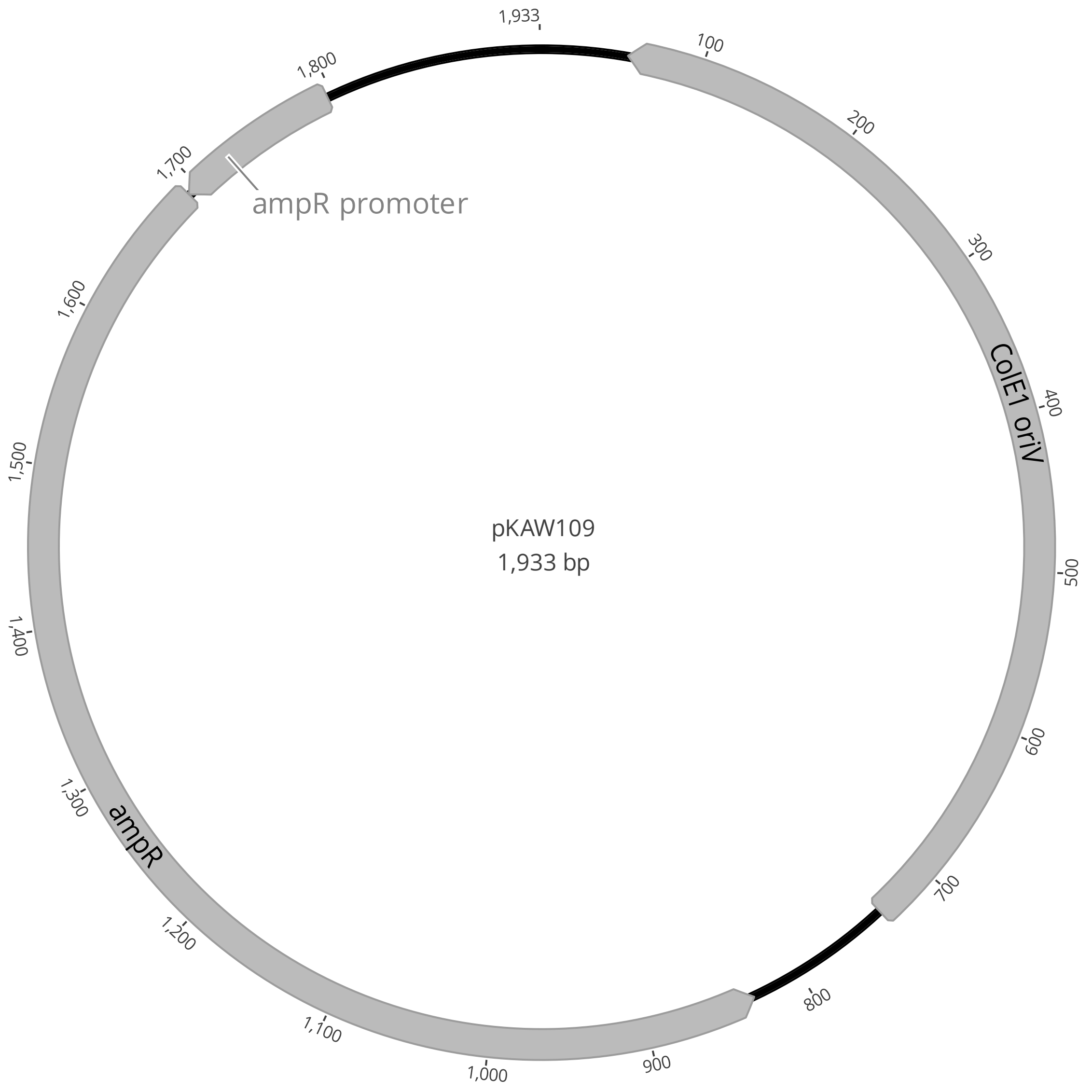


**Figure S5**. pKAW109, the plasmid used in control experiments. The backbone was amplified from pAM041 (Figure S2) but lacks an editing cassette and sgRNA.

**Supplemental Tables**

| **Table S1.** Strains used in this work. All strains except DH5a and DH5a lambda pir contained the helper plasmid pDY118A. Intm designates intermediate strains that had previously been edited and were used for a second round of genome editing. | | | |
|---|---|---|---|
| strain | genotype | notes | Biosample accession |
| *E. coli* DH5α | *Δ(argF-lac)169*, φ80d*lacZ58*(M15), *ΔphoA8*, *glnX44*(AS), *λ*⁻, *deoR481*, *rfbC1*, *gyrA96*(NalR), *recA1*, *endA1*, *thiE1*, *hsdR17* | Coli Genetic Stock Center (CGCS) strain # 14231 | |
| *E. coli* DH5α lambda pir | DH5α with genomic integration of gene encoding Pir | obtained from Eric Stabb (2) | |
| *E. coli* BW25113* | BW25113 (MG1655 *Δ(araD-araB)567*, *ΔlacZ4787*(::rrnB-3), *λ*⁻, *rph-1*, *Δ(rhaD-rhaB)568*, *hsdR514)*; *pdxB* replaced with *kanR* | parental strain for all I-SceI edited strains | SAMN40533614 |
| *E. coli* BW25113* *yifE Q100** | BW25113* with a C → T mutation at position 298 in *yifE* resulting in a Q100* missense mutation | | SAMN40533615 |
| *E. coli* BW25113* *gapA** | BW25113* with a G → T mutation at position 629 in *gapA* resulting in a G210V substitution | | SAMN40533616 |
| *E. coli* BW25113* *gapA* ΔacrZ-pgl* | BW25113* with a G → T mutation at position 629 in *gapA* resulting in a G210V substitution and a 3812 bp deletion spanning genomic positions 790371-794183 | | SAMN40533617 |
| *E. coli* BW25113* *gapA* ΔacrZ-pgl rpoC** | BW25113* with a G → T mutation at position 629 in *gapA* resulting in a G210V substitution, a 3812 bp deletion spanning genomic positions 790371-794183 and a C → A mutation at position 2788 in *rpoC* resulting in an L930M substitution | | SAMN40533618 |
| *E. coli* MG1655* | MG1655 with 82 bp deletion upstream of *pyrE; gmhB / rrsH* intergenic (+240/-123, A → G); *paoD* E266K (GAG → AAG); *sixA* E61A (GAA → GCA)*; gltP / yjcO* intergenic (+587/+55) +GC | parental strain for all Cas9-assisted editing | SAMN40533619 |
| *E. coli* MG1655* *acs*328::mCherry intm A | MG1655*, codons 328-331 of *acs* replaced with mCherry-NGG (CTGTCCCCTCAGTTCATGTAagg); *chaA*/*chaB* intergenic mutation (-120/-150) A → G | unintended mutation between *chaA* and *chaB* occurred while introducing *acs*328::mCherry | SAMN40533620 |
| *E. coli* MG1655* *acs*328::mMCherry intm B | MG1655*, codons 328-331 of *acs* replaced with mCherry-NGG | unintended mutation between *frmR* and *yaiO* | SAMN40533621 |

| | | | |
|---|---|---|---|
| | (CTGTCCCCTCAGTTCATGTAagg), *frmR*/*yaiO* intergenic 1 bp deletion (-132/+56) | occurred while introducing *acs*328::mCherry | |
| *E. coli* MG1655* *acs*328::mCherry intm C | MG1655*, codons 328-331 of *acs* replaced with mCherry-NGG (CTGTCCCCTCAGTTCATGTAagg) | | SAMN40533622 |
| *E. coli* MG1655* *acs*610::mCherry intm | MG1655*, codons 610-624 of *acs* replaced with mCherry-NGG (CTGTCCCCTCAGTTCATGTAagg) | | SAMN40533623 |
| *E. coli* MG1655* NEW_*ackA*105-2::mCherry intm | MG1655*, codons 105-114 of *ackA* replaced with mCherry-NGG (CTGTCCCCTCAGTTCATGTAagg) | | SAMN40533624 |
| *E. coli* MG1655* *adhE*490::mCherry intm A | MG1655*, codons 486-496 of *adhE* replaced with mCherry-NGG (CTGTCCCCTCAGTTCATGTAagg) | | SAMN40533625 |
| *E. coli* MG1655* *adhE*490::mCherry intm B | MG1655*, codons 486-496 of *adhE* replaced with mCherry-NGG (CTGTCCCCTCAGTTCATGTAagg) | | SAMN40533626 |
| *E. coli MG1655** *cyaA*499::mMCherry intm | MG1655*, codons 497-500 of *cya* replaced with mCherry-NGG (CTGTCCCCTCAGTTCATGTAagg) | | SAMN40533627 |
| *E. coli* MG1655* *fbaB*236 syn intm | MG1655*, codons 230-237 of *fbaB* replaced with synonymous codons (ACAATAGGAGcggACATTGTGAAG) | mutated codons include a novel 'cgg' PAM site. | SAMN40533628 |
| *E. coli* MG1655* *pgi*352 syn intm | MG1655*, codons 344-352 of *pgi* replaced with synonymous codons (CGGTTTGCAGCCTATTTTCAACAa) | final codon ending with 'a' is followed by 'gg' in the genome, providing a novel 'agg' PAM site | SAMN40533629 |
| *E. coli* MG1655* *ptsI*513 syn intm | MG1655*, codons 509-514 of *ptsI* replaced with synonymous codons (GAGAGAGCAACCTTGTTACTATTa) | final codon ending with 'a' is followed by 'gg' in the genome, providing a novel 'agg' PAM site | SAMN40533630 |
| *E. coli* MG1655* pykF085/090 syn intm | MG1655*, codons 81-91 of *pykF* replaced with synonymous codons (AATGATGTCAGcctAAAGGCCGGACAAACGTTC) | mutated codons include a reverse complimentary novel 'agg' PAM site | SAMN40533631 |
| *E. coli* MG1655* *pykF*466 syn intm | MG1655* background, codons 451-466 of *pykF* replaced with synonymous codons (ATGGTCAGCGGAGCTTTAGTTCCATCaggAACAACGAATACCGCTAGC) | mutated codons include a reverse complimentary novel 'agg' PAM site | SAMN40533632 |

| **Table S2.** Plasmids used in this work. | | |
|---|---|---|
| plasmid | use / description | source or reference |
| pDY118A (Fig S1) | helper plasmid for Cas9-assisted editing; encodes Cas9 under control of the Tet promoter, lambda Red enzymes under control of a heat-inducible promoter, I-Sce-I under control of the lacIq promoter, and chloramphenicol resistance; pSC101 temperature-sensitive origin of replication | (3) |
| pAM041 (Fig S2) | backbone for donor and guide plasmids used for Cas9-assisted editing (Addgene ID 217969) | this study |
| Cas9 donor plasmids | encode ampicillin resistance, an sgRNA and an editing cassette | |
| pDY339 | sgRNA targets an mCherry-AGG sequence introduced at codon 105 of *ackA* through a previous round of genome editing; repair template introduces three synonymous mutations at codons 105, 113 and 114 (Table S4, GB_ackA105_2nd) | (3) |
| pDY348 | sgRNA targets an mCherry-AGG sequence introduced at codon 499 of *cyaA* through a previous round of genome editing; repair template introduces two synonymous mutations at codons 497 and 498 (Table S4, GB_cyaA499_2nd) | this study |
| pDY449 | sgRNA targets a sequence of synonymous codons introduced at codons 84-92 of *pykF* by a previous round of genome editing; repair template introduces two synonymous mutations at codons 85 and 86 (Table S4, GB_pykF085_2nd) | (3) |
| pDY450 | sgRNA targets a sequence of synonymous codons introduced at codons 84-92 of *pykF* by a previous round of genome editing; repair template introduces five synonymous mutations at codons 83, 84 and 90 (Table S4, GB_pykF090_2nd) | (3) |
| pDY452 | sgRNA targets a sequence of synonymous codons introduced at codons 453-459 of *pykF* by a previous round of genome editing; repair template introduces seven synonymous mutations at codons 452, 453 and 466 (Table S4, GB_pykF466_2nd) | (3) |
| pDY453 | sgRNA targets a sequence of synonymous codons introduced at codons 453-459 of *pykF* by a previous round of genome editing; repair template introduces four synonymous mutations at codons 452 and 453 (Table S4, GB_pykF470_2nd) | (3) |
| Cas9 guide plasmids (Fig S2) | encode ampicillin resistance and an sgRNA but no editing cassette | |
| pDY320 | sgRNA targets an mCherry-AGG sequence introduced at codon 328 of *acs*, codon 610 of *acs* or codon 490 of *adhE* by previous rounds of genome editing | (3) |
| pDY332 | sgRNA targets a sequence of synonymous codons introduced at codons 344-352 of *pgi* by a previous round of genome editing | (3) |
| pDY334 | sgRNA targets a sequence of synonymous codons introduced at codons 230-237 of *fbaB* by a previous round of genome editing | (3) |
| pDY337 | sgRNA targets a sequence of synonymous codons introduced at codon 509-514 of *ptsI* by a previous round of genome editing | this study |
| lambda Red recombineering | encodes GFP and contains an editing casette to replace *pdxB* with *kanR* | |

| | | |
|---|---|---|
| mutation cassette plasmid (Fig S3) | | |
| pKAW086 | donor plasmid for lambda Red recombineering; editing cassette contains *kan* with homology arms flanking *E. coli pdxB*; encodes GFP under control of a constitutive promoter; R6K origin of replication that requires Pir to be maintained in cells; kanamycin and ampicillin resistance (Addgene ID 217967) | this study |
| I-SceI mutation cassette plasmid (Fig S4) | editing cassette used to introduce a point mutation in *rpoS* via I-SceI-assisted editing; ampicillin resistance | |
| pKAW066 | donor plasmid for I-SceI assisted editing; editing cassette comprised of three segments: first, a segment containing a homology arm (HA) upstream of the edit, the edit and a short downstream HA; second, a segment containing an I-SceI cut-site and a streptomycin resistance gene; and third, a final segment containing a short HA upstream of the edit, the edit and a downstream HA; ampicillin resistance; pMB1 origin of replication (Addgene ID 217968) | this study |
| Control plasmid (Plasmid lacking editing cassette and sgRNA) (Fig S5) | ampicillin resistance | |
| pKAW109 | empty plasmid used for Cas9-assisted editing control experiments; ampicillin resistance; pMB1 origin of replication | this study |

| **Table S3.** PCR primers used in this work. F and R designate forward and reverse primers, respectively. | | |
|---|---|---|
| *Primers for amplification of the donor and guide plasmid backbones used in Cas9-assisted genome editing* | | |
| primer | sequence | use |
| Donor2_bbF | GTTTTAGAGCTAGAAATAGCAAGTTAAAATAAGGCTAGTCCGTTATCAAC | amplification of donor plasmid backbone from pAM041 |
| Donor2_bbR | ACAGCTTCAAGTAGTCGGGGATGTC | |
| P623_RmvSpacerF | GTTTTAGAGCTAGAAATAGC | amplification of guide plasmid backbone from pAM041 |
| P624_ RmvSpacerR | ACTAGTATTATACCTAGGAC | |
| *Primer for Sanger sequencing donor and guide plasmid inserts* | | |
| primer | sequence | use |
| 2nd_seqF | ATAAGGGCGACACGGAAATGTTGAATACTC | sequencing primer to verify insert in donor and guide plasmids |
| *Primers for colony PCR and/or Sanger sequencing to confirm Cas9-assisted editing* | | |
| primer | sequence | target gene / amplifies region around indicated edited codon(s) |
| acs328-SeqF | GCATGTGGTGGTACTGAAGCGTACTG | *acs* / codon 328 |
| acs328-SeqR | CTTCCGGGTTAATTGGCTCGCCCAC | *acs* / codon 328 |
| acs610-SeqF | GGAACCGTCACCAGAACTGTACGCAGAAG | *acs* / codon 610 |
| acs610-SeqR | CAAACCGTTACCGACTCGCATCGGGC | *acs* / codon 610 |
| adhE490_seq_F | TCTCAGGGTGGTATCGGTGACCTG | *adhE* / codon 490 |
| adhE490_seq_R | TGTTGTGGAAGCCGTTATAGTGCCTCAG | *adhE* / codon 490 |
| cyaA499_seq_F | GCTGGAAGCCTATTCCTGGGAATAC | *cyaA* / codon 499 |
| cyaA499_seq_R | CAGTTTCTGTACCGATACATTCAGGCG | *cyaA* / codon 499 |
| fbaB238_seq_F | CTGTATGCCAGCGTGGAGCAGG | *fbaB* / codon 238 |
| fbaB238_seq_R | CTGCACGGCGTTAATCAGTTTCACG | *fbaB* / codon 238 |
| pgi352_seq_F | CGTTGGCGAGTTTGGTATTG | *pgi* / codon 352 |
| pgi352_seq_R | GATGATCAGAGAGCGGGTTATG | *pgi* / codon 352 |
| ptsI_3'F | GAGCTGGAGCAGGAAATCATAG | *ptsI* / codon 513 |
| ptsI_3'R | GAGAATGCGTGGTTGGTTTC | |
| pykF_seq_F | GAACTTCTCTCATGGTGACTATGCAG | *pykF* / codons 85 and 466 |
| pykF_seq_R | GATGCTTCCATCGGATTCATCTTAGATA | |
| *Primers used for lambda Red recombineering* | | |
| primer | sequence | use |
| KW459F | CTCCCTGACCTGGTGGTTGC | amplification of *pdxB* editing cassette from pKAW086 |
| KW460R | ACTGACGTTCAGCCAGCGTTTCAA | |
| KW438F | AAAATATGCCTTATGCCCGCGACTT | bind within *pdxB* coding sequence to check for presence of *pdxB* in edited strains |
| KW439R | GCGTTAAAACCCAGTTTACACAGCAATGAT | |

| | | |
|---|---|---|
| KW440F | GCGTCAGTGAGGTCACTTTCGT | bind outside of *pdxB* coding sequence to check for correct insertion of *kanR* |
| KW441R | ATCCTGCACGGTGATTGTCTTACC | |
| *Primers used for I-SceI-assisted editing* | | |
| primer | sequence | use |
| KW061F | CACTTGGTTCATGGTCCAGCTTA | amplification of *rpoS* editing cassette from pKAW066 that inserts a T at position 98 in *rpoS* |
| KW062R | CAGTACATCAACCAGTACGCCTA | |
| KW385F | TAGCACCGGAACCAGTTCAACAC | bind outside of *rpoS* coding sequence; used to confirm correct editing |
| KW384R | GTTGTCGGTAGCAGACGGTCTGA | |
| *Primers used for pKAW109 construction* | | |
| primer | sequence | use |
| KW027F | CGCAGGAAAGAACATGTG | amplification of oriV and ampicillin resistance gene from pAM041 |
| KW181R | CCAGAAATCATCCTTAGCGAA | |

**Table S4.** gBlocks used to construct donor plasmids used for Cas9-assisted editing.

| Lower-case italics, extensions to direct Gibson Assembly with vector backbone; upper-case black, editing cassette; green, J23119 constitutive promoter; blue, 20-nt gRNA spacer sequence; red and magenta, target and synonymous immunizing mutations in or near the PAM site, respectively. (Target codons are underlined.) | |
|---|---|
| GB_ackA105_2nd (for construction of pDY339) | *caaggcctacgtgaagcaccccgccgacatccccgactacttgaagctgt*GGCTTTAGGTGCAGGCGCCGCTCACAGCGAAGCGCTCAACTTTATCGTTAATACTATTCTGGCACAAAAACCAGAACTGTCTGCGCAGCTGACTGCTATCGGTCACCGTATCGTACACGGCGGCGAAAAGTATACCAGCTCCGTAGTGATAGATGAGTCTGTTATTCAGGGTATTAAGGATGCAGCTTCTTTTGCACCGCTGCACAACCCGGCTCACCTGATCGGTATCGAAGAAGCTCTGAAATCTTTCCCACAGCTGAAAGACAAAAACGTTGCTGTATTTGACACCGCGTTCCACCAGACTATGCCGGAAGAGTCTTACCTCTTTGACAGCTAGCTCAGTCCTAGGTATAATACTAGTCTGTCCCCTCAGTTCATGTA*gttttagagctagaaatagcaagttaaaataaggctagtccgttatcaac* |
| GB_cyaA499_2nd (for construction of pDY348) | *caaggcctacgtgaagcaccccgccgacatccccgactacttgaagctgt*CCCGATCTCTCGGAACCGAATCTGACCTTTATTTATGTGCCGCCGGGCCGGGCTAACCGTTCAGGTTGGTATCTGTATAACCGCGCGCCAAATATTGAGTCGATCATCAGCCATCAGCCGCTGGAATATAACCGTTACCTGAATAAACTGGTGGCGTGGGCATGGTTCAATGGCCTGCTGACCTCGCGCACCCGTTTGTATATTAAAGGTAACGGCATTGTCGATTTGCCTAAGTTGCAGGAGATGGTCGCCGACGTGTCGCACCATTTCCCGCTGCGCTTACCTGCACCGACACCGAAGGCGCTCTACAGCCCGTGTGAGTTGACAGCTAGCTCAGTCCTAGGTATAATACTAGTCTGTCCCCTCAGTTCATGTA*gttttagagctagaaatagcaagttaaaataaggctagtccgttatcaac* |
| GB_pykF085_2nd (for construction of pDY449) | *caaggcctacgtgaagcaccccgccgacatccccgactacttgaagctgt*GAACTTCTCTCATGGTGACTATGCAGAACACGGTCAGCGCATTCAGAATCTGCGCAACGTGATGAGCAAAACTGGTAAAACCGCCGCTATCCTGCTTGATACCAAAGGTCCGGAAATCCGCACCATGAAACTGGAAGGCGGTAACGACGTTTCTTTGAAGGCTGGTCAGACCTTTACTTTCACCACTGATAAATCTGTTATCGGCAACAGCGAAATGGTTGCGGTAACGTATGAAGGTTTCACTACTGACCTGTCTGTTGGCAACACCGTACTGGTTGACGATGGTCTGATCGGTATGGAAGTTACCGCCTTGACAGCTAGCTCAGTCCTAGGTATAATACTAGTTGAACGTTTGTCCGGCCTTT*gttttagagctagaaatagcaagttaaaataaggctagtccgttatcaac* |
| GB_pykF090_2nd (for construction of pDY450) | *caaggcctacgtgaagcaccccgccgacatccccgactacttgaagctgt*GAACTTCTCTCATGGTGACTATGCAGAACACGGTCAGCGCATTCAGAATCTGCGCAACGTGATGAGCAAAACTGGTAAAACCGCCGCTATCCTGCTTGATACCAAAGGTCCGGAAATCCGCACCATGAAACTGGAAGGCGGTAACGACGTGAGCCTGAAAGCTGGTCAGACGTTTACTTTCACCACTGATAAATCTGTTATCGGCAACAGCGAAATGGTTGCGGTAACGTATGAAGGTTTCACTACTGACCTGTCTGTTGGCAACACCGTACTGGTTGACGATGGTCTGATCGGTATGGAAGTTACCGCCATTGAAGGTAACTTGACAGCTAGCTCAGTCCTAGGTATAATACTAGTTGAACGTTTGTCCGGCCTTT*gttttagagctagaaatagcaagttaaaataaggctagtccgttatcaac* |

| | |
|---|---|
| GB_pykF466_2nd (for construction of pDY452) | *caaggcctacgtgaagcaccccgccgacatccccgactacttgaagctgt*CGAAAAAACGGCTC ATCAGTTGGTACTGAGCAAAGGCGTTGTGCCGCAGCTTGTTAAAGAG ATCACTTCTACTGATGATTTCTACCGTCTGGGTAAAGAACTGGCTCTGC AGAGCGGTCTGGCACACAAAGGTGACGTTGTAGTTATGGTGAGCGGT GCACTGGTACCGAGCGGCACTACTAACACCGCAAGCGTTCACGTCCT GTAATATTGCTTTTGTGAATTAATTTGTATATCGAAGCGCCCTGATGGGC GCTTTTTTTATTTAATCGATAACCAGAAGCAATAAAAAATCAAATCGGA TTTCACTATATAATCTCACTTTATCTAAGATGAATCCGATTTGACAGCTA GCTCAGTCCTAGGTATAATACTAGTAGCGGAGCTTTAGTTCCATC*gttttag agctagaaatagcaagttaaaataaggctagtccgttatcaac* |
| GB_pykF470_2nd (for construction of pDY453) | *caaggcctacgtgaagcaccccgccgacatccccgactacttgaagctgt*CGAAAAAACGGCTC ATCAGTTGGTACTGAGCAAAGGCGTTGTGCCGCAGCTTGTTAAAGAG ATCACTTCTACTGATGATTTCTACCGTCTGGGTAAAGAACTGGCTCTGC AGAGCGGTCTGGCACACAAAGGTGACGTTGTAGTTATGGTGAGCGGT GCACTGGTACCGAGCGGCACTACTAACACCGCATCTGTTCACGTCCTG TAATATTGCTTTTGTGAATTAATTTGTATATCGAAGCGCCCTGATGGGCG CTTTTTTTATTTAATCGATAACCAGAAGCAATAAAAAATCAAATCGGAT TTCACTATATAATCTCACTTTATCTAAGATGAATCCGATTTGACAGCTAG CTCAGTCCTAGGTATAATACTAGTAGCGGAGCTTTAGTTCCATC*gttttagag ctagaaatagcaagttaaaataaggctagtccgttatcaac* |

**Table S5**. 120-nt ssDNA editing cassettes used for Cas9-assisted editing and the associated spacer sequences used to direct Cas9 genome cleavage. Red and magenta, target and immunizing mutations, respectively. Target codons are underlined.

| target gene/codon | sequence | associated spacer sequence |
|---|---|---|
| *acs* / 328 | GTGACCGGACACAGTTACTTGCTGTACGGCCCGCTGGCCTG CGGTGCGACCACGCTCATGTTCGAGGGCGTACCCAACTGG CCGACGCCTGCCCGTATGGCGCAGGTGGTGGACAAGCAT | CTGTCCCCTC AGTTCATGTA |
| *acs* / 328 | GTGACCGGACACAGTTACTTGCTGTACGGCCCGCTGGCCTG CGGTGCGACCACGCTCATGTTTGAAGGCGTACCCAACTGGC CGACGCCTGCCCGTATGGCGCAGGTGGTGGACAAGCAT | CTGTCCCCTC AGTTCATGTA |
| *acs* / 328 | GTGACCGGACACAGTTACTTGCTGTACGGCCCGCTGGCCTG CGGTGCGACCACGCTGATGTTCGAAGGCGTACCCAACTGG CCGACGCCTGCCCGTATGGCGCAGGTGGTGGACAAGCAT | CTGTCCCCTC AGTTCATGTA |
| *acs* / 328 | GTGACCGGACACAGTTACTTGCTGTACGGCCCGCTGGCCTG CGGTGCGACCACGCTGATGTTTGAGGGCGTACCCAACTGG CCGACGCCTGCCCGTATGGCGCAGGTGGTGGACAAGCAT | CTGTCCCCTC AGTTCATGTA |
| *acs* / 610 | GACGTGCTGCACTGGACCGACTCCCTGCCTAAAACCCGCTC CGGCAAAATCATGCGCCGTATTCTGCGTAAGATTGCGGCGG GCGATACCAGCAACCTGGGCGATACCTCGACGCTTGCC | CTGTCCCCTC AGTTCATGTA |
| *acs* / 619 | TGGACCGACTCCCTGCCTAAAACCCGCTCCGGCAAAATTAT GCGCCGTATTCTGCGTAAGATTGCAGCGGGCGATACCAGCA ACCTGGGCGATACCTCGACGCTTGCCGATCCTGGCGTA | CTGTCCCCTC AGTTCATGTA |
| *acs* / 624 | ACTCCCTGCCTAAAACCCGCTCCGGCAAAATTATGCGCCGT ATTCTGCGTAAGATTGCGGCGGGCGATACCTCGAACCTGGG CGATACCTCGACGCTTGCCGATCCTGGCGTAGTCGAG | CTGTCCCCTC AGTTCATGTA |
| *acs* / 610 | GACGTGCTGCACTGGACCGACTCCCTGCCTAAAACCCGCTC CGGCAAAATCATGCGCCGTATTCTGCGCAAAATTGCGGCGG GCGATACCAGCAACCTGGGCGATACCTCGACGCTTGCC | CTGTCCCCTC AGTTCATGTA |
| *acs* / 610 | GACGTGCTGCACTGGACCGACTCCCTGCCTAAAACCCGCTC CGGCAAAATTATGCGCCGTATTCTGCGTAAAATTGCGGCGG GCGATACCAGCAACCTGGGCGATACCTCGACGCTTGCC | CTGTCCCCTC AGTTCATGTA |
| *acs* / 610 | GACGTGCTGCACTGGACCGACTCCCTGCCTAAAACCCGCTC CGGCAAAATTATGCGCCGTATTCTGCGCAAGATTGCGGCGG GCGATACCAGCAACCTGGGCGATACCTCGACGCTTGCC | CTGTCCCCTC AGTTCATGTA |
| *pgi* / 352 | CACAACGTTACCGTTACGGTCAACATACTTACCGTTGGACT CCATATTCCCCTGCTGGAAGTACGCCGCAAACCGGTGCATA TACTGGTCATACGGCAGAATCGCTTCAGTTTCCGCACC | TTTGCAGCCT ATTTTCAACA |
| *pgi* / 352 | CACAACGTTACCGTTACGGTCAACATACTTACCGTTGGACT CCATATTGCCCTGCTGGAAGTACGCCGCGAACCGGTGCATA TACTGGTCATACGGCAGAATCGCTTCAGTTTCCGCACC | TTTGCAGCCT ATTTTCAACA |
| *pgi* / 352 | CACAACGTTACCGTTACGGTCAACATACTTACCGTTGGACT CCATATTGCCCTGCTGGAAGTACGCCGCAAAACGGTGCATA TACTGGTCATACGGCAGAATCGCTTCAGTTTCCGCACC | TTTGCAGCCT ATTTTCAACA |
| *pgi* / 352 | CACAACGTTACCGTTACGGTCAACATACTTACCGTTGGACT CCATATTCCCCTGCTGGAAGTACGCCGCGAAACGGTGCATA TACTGGTCATACGGCAGAATCGCTTCAGTTTCCGCACC | TTTGCAGCCT ATTTTCAACA |
| *fbaB* / 238 | TGACCGGTCAGGCAAACCATCTGGCGGCAACCATCGGTGC AGATATCGTCAAACAAAAAATGGCCGAGAATAACGGCGGC TATAAAGCAATTAATTACGGTTACACCGACGATCGTGTTT | TCTGGCGGCA ACAATAGGAG |

| | | |
|---|---|---|
| *adhE* / 486 | CGCTGGATGAAGTGATTACTGATGGCCACAAACGTGCGCTC ATCGTGACCGACCGGTTTCTGTTCAACAATGGTTATGCTGAT CAGATCACTTCCGTACTGAAAGCAGCAGGCGTT | CTGTCCCCTC AGTTCATGTA |
| *adhE* / 490 | CGCTGGATGAAGTGATTACTGATGGCCACAAACGTGCGCTC ATCGTGACTGACCGGTTTTTATTCAACAATGGTTATGCTGAT CAGATCACTTCCGTACTGAAAGCAGCAGGCGTTGAAA | CTGTCCCCTC AGTTCATGTA |
| *adhE* / 496 | AAGTGATTACTGATGGCCACAAACGTGCGCTCATCGTGACT GACCGGTTTCTGTTCAACAATGGTTATGCTGATCAGATCACT TCCGTACTGAAAGCAGCAGGCGTTGAAACTGAAGTCT | CTGTCCCCTC AGTTCATGTA |
| *adhE* / 496 | AAGTGATTACTGATGGCCACAAACGTGCGCTCATCGTGACT GACCGGTTTCTGTTCAACAATGGTTATGCCGATCAGATCACT TCCGTACTGAAAGCAGCAGGCGTTGAAACTGAAGTCT | CTGTCCCCTC AGTTCATGTA |
| *adhE* / 496 | AAGTGATTACTGATGGCCACAAACGTGCGCTCATCGTGACT GACCGGTTTCTGTTCAACAATGGTTATGCGGATCAGATCAC TTCCGTACTGAAAGCAGCAGGCGTTGAAACTGAAGTCT | CTGTCCCCTC AGTTCATGTA |

| **Table S6.** Lambda Red recombineering editing cassette used to replace *pdxB* with *kanR* |
|---|
| Red, homology arm (HA) upstream of *pdxB*; black, sequences containing promoters, terminators and FRT sites; grey, *kanR*; cyan, HA downstream of *pdxB*. |
| CTCCCTGACCTGGTGGTTGCCCAGGAGGAGGGCCGGAAATAGGTTGTATCATTACGTATC<br>CTTATACCTGAAATCTTCGCAAGTATGCCTGGCCGCGAGATTATGGCACACTTGTCCGGTT<br>AACTCTCGTCTCATACAGGTAACACAAACGTGATTCCGGGGATCCGTCGACCTGCAGTTCG<br>AAGTTCCTATTCTCTAGAAAGTATAGGAACTTCAGAGCGCTTTTGAAGCTCACGCTGCCGC<br>AAGCACTCAGGGCGCAAGGGCTGCTAAAGGAAGCGGAACACGTAGAAAGCCAGTCCGCA<br>GAAACGGTGCTGACCCCGGATGAATGTCAGCTACTGGGCTATCTGGACAAGGGAAAACGC<br>AAGCGCAAAGAGAAAGCAGGTAGCTTGCAGTGGGCTTACATGGCGATAGCTAGACTGGG<br>CGGTTTTATGGACAGCAAGCGAACCGGAATTGCCAGCTGGGGCGCCCTCTGGTAAGGTTG<br>GGAAGCCCTGCAAAGTAAACTGGATGGCTTTCTTGCCGCCAAGGATCTGATGGCGCAGGG<br>GATCAAGATCTGATCAAGAGACAGGATGAGGATCGTTTCGCATGATTGAACAAGATGGAT<br>TGCACGCAGGTTCTCCGGCCGCTTGGGTGGAGAGGCTATTCGGCTATGACTGGGCACAAC<br>AGACAATCGGCTGCTCTGATGCCGCCGTGTTCCGGCTGTCAGCGCAGGGGCGCCCGGTTCT<br>TTTTGTCAAGACCGACCTGTCCGGTGCCCTGAATGAACTGCAGGACGAGGCAACGCGGCT<br>ATCGTGGCTGGCCACGACGGGCGTTCCTTGCGCAGCTGTGCTCGACGTTGTCACTGAAGCG<br>GGAAGGGACTGGCTGCTATTGGGCGAAGTGCCGGGGCAGGATCTCCTGTCATCTCACCTT<br>GCTCCTGCCGAGAAAGTATCCATCATGGCTGATGCAATGCGGCGGCTGCATACGCTTGAT<br>CCGGCTACCTGCCCATTCGACCACCAAGCGAAACATCGCATCGAGCGAGCACGTACTCGG<br>ATGGAAGCCGGTCTTGTCGATCAGGATGATCTGGACGAAGAGCATCAGGGGCTCGCGCCA<br>GCCGAACTGTTCGCCAGGCTCAAGGCGCGCATGCCCGACGGCGAGGATCTCGTCGTGACC<br>CATGGCGATGCCTGCTTGCCGAATATCATGGTGGAAAATGGCCGCTTTTCTGGATTCATCG<br>ACTGTGGCCGGCTGGGTGTGGCGGACCGCTATCAGGACATAGCGTTGGCTACCCGTGATA<br>TTGCTGAAGAGCTTGGCGGCGAATGGGCTGACCGCTTCCTCGTGCTTTACGGTATCGCCGC<br>TCCCGATTCGCAGCGCATCGCCTTCTATCGCCTTCTTGACGAGTTCTTCTAATAAGGGGAT<br>CTTGAAGTTCCTATTCCGAAGTTCCTATTCTCTAGAAAGTATAGGAACTTCGAAGCAGCTC<br>CAGCCTACAGTGCATCATCCGGCACGTTAATCTCTTCTTCATGCTCTCTGCTGTAACATTGG<br>CAGGGAGCTTTGCTATTTCTGGAGTAAACCACCATGTCTGAAGGCTGGAACATTGCCGTCC<br>TGGGCGCAACTGGCGCTGTGGGCGAAGCCCTGCTTGAAACGCTGGCTGAACGTCAGT |

| **Table S7.** Editing cassette used to insert T at position 98 in *rpoS* with I-SceI-assisted editing |
| --- |
| Red, homology arm (HA) upstream of the edit; yellow, edit; cyan, downstream HA; black, segment containing terminators; green, segment containing an I-SceI cut-site; black, segment containing a promoter; grey, streptomycin resistance gene; red, short HA upstream of the edit; yellow, edit; blue, downstream HA. |
| CACTTGGTTCATGGTCCAGCTTATGGGACAACTCACGTGCGGTTCGCAGGTAAACGTTCAG<br>CTCCTTTACGATGTGAATCGGCAAACGAATAGTACGGGTTTGGTTCATAATCGCCCGTTCA<br>ATCGTCTGGCGAATCCACCAGGTTGCGTATGTTGAGAAGCGGAAACCACGTTCCGGGTCA<br>AACTTCTCTACCGCGCGGATCAGCCCCAGGTTGCCCTCTTCGATAAGGTCCAGCAACGCCA<br>GACCACGATTGCCATAACGGCGGGCAATTTTTACCACCAGACGCAAGTTACTCTCGATCAT<br>CCGGCGGCGAGAGGCGACATCTCCACGCAGTGCGCGACGCGCAAAATAAACTTCTTCTTC<br>GGCCGTTAACAGTGGTGAATAACCAATCTCACCAAGGTAAAGCTGAGTCGCGTCCAACAC<br>ACGCTGTGTGGCTCCCTGCGATAACAGTTCCTCTTCGGCCAAATCGTTATCACTGGGTTCC<br>TTGTTCTACTAAGGCCATCTCAAGAGTGGCAGCGGTTCTGTTAAGTAACTGAACCCAATGT<br>CGTTAGTGACGCTTACCCGCAAAAAACCCCGCTTCGGCGGGGTTTTTCGCTCTTAAGAGG<br>TCACTGACCTAACAAAAAAAAAACCCCGCCCCTGACAGGGCGGGGTTTTTTTGGTCTTGA<br>GTGGCAGAGTCAGTTATCGCGAGCAGTATGTAAGTAGATCCTCAGTGTCAGCTAGGGATA<br>ACAGGGTAATCCTGGTGTCCCTGTTGATACCGGGAAGCCCTGGGCCAACTTTTGGCGAAA<br>ATGAGACGTTGATCGGCACGTAAGAGGTTCCAACTTTCACCATAATGAAATAAGATCACT<br>ACCGGGCGTATTTTTGAGTTATCGAGATTTCAGGAGCTAAGGAAGCTAAAATGAGGGA<br>AGCGGTGATCGCCGAAGTATCGACTCAACTATCAGAGGTAGTTGGCGTCATCGAGCGCCA<br>TCTCGAACCGACGTTGCTGGCCGTACATTTGTACGGCTCCGCAGTGGATGGCGGCCTGAA<br>GCCACACAGTGATATTGATTTGCTGGTTACGGTGACCGTAAGGCTTGATGAAACAACGCG<br>GCGAGCTTTGATCAACGACCTTTTGGAAACTTCGGCTTCCCCTGGAGAGAGCGAGATTCTC<br>CGCGCTGTAGAAGTCACCATTGTTGTGCACGACGACATCATTCCGTGGCGTTATCCAGCTA<br>AGCGCGAACTGCAATTTGGAGAATGGCAGCGCAATGACATTCTTGCAGGTATCTTCGAGC<br>CAGCCACGATCGACATTGATCTGGCTATCTTGCTGACAAAAGCAAGAGAACATAGCGTTG<br>CCTTGGTAGGTCCAGCGGCGGAGGAACTCTTTGATCCGGTTCCTGAACAGGATCTATTTGA<br>GGCGCTAAATGAAACCTTAACGCTATGGAACTCGCCGCCCGACTGGGCTGGCGATGAGCG<br>AAATGTAGTGCTTACGTTGTCCCGCATTTGGTACAGCGCAGTAACCGGCAAAATCGCGCC<br>GAAGGATGTCGCTGCCGACTGGGCAATGGAGCGCCTGCCGGCCCAGTATCAGCCCGTCAT<br>ACTTGAAGCTAGACAGGCTTATCTTGGACAAGAAGAAGATCGCTTGGCCTCGCGCGCAGA<br>TCAGTTGGAAGAATTTGTCCACTACGTGAAAGGCGAGATCACCAAGGTAGTCGGCAAATA<br>ATTATCACTGGGTTCCTTGTTCTACTAAGGCCTTTTCGTCAAAAACCTCAACTCCGTTCTCA<br>TCAAATTCCGCATCTTCATTTAAATCATGAACTTTCAGCGTATTCTGACTCATAAGGTGGC<br>TCCTACCCGTGATCCCTTGACGGAACATTCAAGCAAAAGCCTGGTTCCGCCGATTTATCGC<br>TGCGGCAAATAACGCAGCGGGTTTACGGATTTCCCCTTGTAACGAATTTCAAAATGCAAG<br>CGTGTTGAACTGGTTCCGGTGCTACCCATGGTCGCTATTTTTTGCCCCGCCTTAACTTCTTG<br>TTGTTCCCGGACCAGCATTGTGTCGTTATGGGCGTAGGCACTCAGGTAATCATCATTATGT<br>TTGATGATAATCAGATTACCGTAGCCGCGCAGCGCGTTACCAGCATAAACAACGCGGCCA<br>TCTGCGGTCGCGATAATTGCCTGTCCTTTGCTGCCTGCGATATCAATCCCCTTGTTGCCCCC<br>CTCAGAAGCGCCAAAGGTTTCGATCACTTTGCCCTCAGTCGGCCAGCGCCAGGTGGAGAT<br>AGGCGTACTGGTTGATGTACTG |

| **Table S8.** 60-nt oligonucleotides containing spacer sequences used to make guide plasmids | | |
|---|---|---|
| Uppercase, 20-nt spacer; lowercase, sequences that overlap the guide plasmid backbone. The complementary 60-nt oligonucleotide fragments were annealed and then ligated into the pAM041 backbone by Gibson assembly. | | |
| primer | sequence | guide plasmid constructed |
| mCherry_sgRNA_F | gtcctaggtataatactagtCTGTCCCCTCAGTTCATGTAgt tttagagctagaaatagc | pDY320 |
| mCherry_sgRNA_R | gctatttctagctctaaaacTACATGAACTGAGGGGACAG actagtattatacctaggac | pDY320 |
| pgi352_2nd_sgRNA_F | gtcctaggtataatactagtTTTGCAGCCTATTTTCAACAgt tttagagctagaaatagc | pDY332 |
| pgi352_2nd_sgRNA_R | gctatttctagctctaaaacTGTTGAAAATAGGCTGCAAAa ctagtattatacctaggac | pDY332 |
| fbaB_2nd_sgRNA_F | gtcctaggtataatactagtTCTGGCGGCAACAATAGGAG gttttagagctagaaatagc | pDY334 |
| fbaB_2nd_sgRNA_R | gctatttctagctctaaaacCTCCTATTGTTGCCGCCAGAa ctagtattatacctaggac | pDY334 |
| ptsI_2nd_sgRNA_F | gtcctaggtataatactagtAGAGCAACCTTGTTACTATTg ttttagagctagaaatagc | pDY337 |
| ptsI_2nd_sgRNA_R | gctatttctagctctaaaacAATAGTAACAAGGTTGCTCTa ctagtattatacctaggac | pDY337 |
